# A multi-omics integrative approach unravels novel genes and pathways associated with senescence escape after targeted therapy in NRAS mutant melanoma

**DOI:** 10.1101/2023.02.13.528114

**Authors:** Vincent Gureghian, Hailee Herbst, Ines Kozar, Katarina Mihajlovic, Noël Malod-Dognin, Gaia Ceddia, Cristian Angeli, Christiane Margue, Tijana Randic, Demetra Philippidou, Milène Tetsi Nomigni, Ahmed Hemedan, Leon-Charles Tranchevent, Joseph Longworth, Mark Bauer, Apurva Badkas, Anthoula Gaigneaux, Arnaud Muller, Marek Ostaszewski, Fabrice Tolle, Nataša Pržulj, Stephanie Kreis

**Author notes:** co-last author (Nataša Pržulj, Stephanie Kreis). **Competing interests:** The authors declare that they have no competing interests.

## Abstract

Therapy Induced Senescence (TIS) leads to sustained growth arrest of cancer cells. The associated cytostasis has been shown to be reversible and cells escaping senescence further enhance the aggressiveness of cancers. Together with targeted therapeutics, senolytics, specifically targeting senescent cancer cells, constitute a promising avenue for improved cancer treatments. Understanding how cancer cells evade senescence is needed to optimise the clinical benefits of this therapeutic approach. Here we characterised the response of three different NRAS mutant melanoma cell lines to a combination of CDK4/6 and MEK inhibitors over 33 days. Transcriptomic data show that all cell lines trigger a senescence programme coupled with strong induction of interferons. Kinome profiling revealed the activation of Receptor Tyrosine Kinases (RTKs) and enriched downstream signaling of neurotrophin, ErbB and insulin pathways. Characterisation of the miRNA interactome associates miR-211-5p with resistant phenotypes. Finally, iCELL-based integration of bulk and single-cell RNA-seq data identified biological processes perturbed during senescence, and predicts new genes involved in its escape. Overall, our data associate insulin signaling with persistence of a senescent phenotype and suggest a new role for interferon gamma in senescence escape through the induction of EMT and the activation of ERK5 signaling.

## INTRODUCTION

Hayflick first observed in 1961 that fibroblasts stop proliferating after 50 passages and display an enlarged and flattened phenotype characteristic of senescence [1, 2]. This observation was later explained by the progressive shortening of telomeres upon replication, which ultimately triggers a DNA Damage Response (DDR) leading to cell cycle arrest [3, 4]. This process was called “replicative senescence”. Several cellular events and environmental conditions induce senescence, and it now emerges as a generic stress response involved in central biological processes such as embryonic development, wound healing and aging [2, 4, 5]. Many stressors that induce senescence (*e.g.*, oxidative stress, radiation), induce DNA damage and activate DDR. This leads to engagement of the Senescence Associated Secretory Phenotype (SASP), one of the characteristic features of senescence in which cells produce and secrete cytokines and growth factors. The SASP itself can trigger senescence in adjacent cells and contributes to the emergence of an inflammatory environment as the senescent cells, which are not eliminated by the immune system, accumulate in the organism [5]. This accumulation leads to the development of degenerative and hyperplastic pathologies and carcinogenesis, linking senescence to aging [6, 7]. Research in this field is challenging due to the heterogeneous and dynamic nature of the SASP and the lack of a universal marker for senescence.

Focusing on cancer, the activation of known oncogenes results in a constant proliferative signaling which causes replication forks to trigger DNA damage and drive primary cells to undergo oncogene-induced senescence (OIS) [8]. Therefore, senescence may be considered as an early evolutionary mechanism of protection against cancer leading to cell cycle arrest of over-proliferating cells [9]. However, cells can escape this arrest and re-establish growth, ultimately leading to the development of cancer. Cancer cells can also undergo senescence when exposed to therapies, so called Therapy-induced senescence (TIS).

Thus, senescence appears as a natural barrier against cancer, but recent works suggest that senescence may be a reversible process [10]. Exit from senescence has been recently linked with the stem-like properties of cancer cells, reinforcing the idea that aside from the SASP, senescence itself may be a transient state in carcinogenesis.

Melanoma arises from the deregulated proliferation of melanocytes representing the deadliest type of skin cancer accounting for about 75% of related deaths. Standard of care involves targeted therapies and immunotherapies [11]. For the BRAF mutated subtype, representing half of the cases, specific first line inhibitors targeting both MEK and BRAF have been developed. The NRAS mutated subtype, which accounts for a quarter of all melanoma cases, lacks such a targeted approach but ongoing clinical trials are assessing the effects of combined MEK and CDK4/6 inhibitors (NCT01781572; NCT02065063). However, tolerance or resistance to such targeted treatments inevitably develops.

MicroRNAs have been shown to be involved in resistance to treatment. miRNAs are 20-22 nucleotides short RNAs, which in complex with the Argonaute (AGO) protein, act as post-transcriptional regulators of gene expression [12]. Canonically, miRNAs destabilise mRNAs by binding to a complementary sequence within the 3’ UTR, this resulting in downregulation of the mRNA and the encoded protein [13]. Experimental methods based on immune-precipitation and sequencing have been developed to profile miRNA-mRNA interactions. The qCLASH method has the advantage of capturing direct physical interactions only, through an additional intermolecular ligation step linking mRNAs to their bound miRNAs [14].

To characterise the response of 3 NRAS mutant melanoma cell lines to a combination of MEK and CDK4/6 inhibitors, we treated the cells over 33 days and collected samples at multiple time points. Samples were analyzed by total RNA-seq, small RNA-seq, qCLASH and kinome profiling. Our results recapitulate several previous observations while displaying an unforeseen interplay between interferon signalling and senescence. Additionally, we uncover deregulated processes associated with the onset of senescence and its exit by adapting a non-negative matrix tri-factorization (NMTF)-based methodology, “iCell” [15], to integrate bulk RNA-seq and single-cell data. Moreover, the dimensionality reduction and the clustering properties of the NMTF allowed us to associate novel genes to senescence escape.

## RESULTS

### Experimental design and description of the data sets

To study the effects of MEKi and CDK4/6i, 3 NRAS mutant melanoma cell-lines were selected: IPC298, MELJUSO and SKMEL30. We determined the GI50 (drug concentration at which cell growth is reduced by half) and opted for a concentration of 16nM, 110nM and 35nM of Binimetinib (MEKi) for IPC298, MELJUSO and SKMEL30, respectively and 1 *µ*M for Palbociclib (CDK4/6i) as this inhibitor led to incomplete dose-response curves (Supp. Table 1). Upon treatment over 33 days, distinct cellular responses (Fig. 1) were observed. IPC298 suffered very little from the treatment showing constant proliferation and no morphological change. SKMEL30 stopped proliferating and accumulated pigmentation before restoring proliferation after around 14 days. Finally, MELJUSO stopped dividing and started to stretch to acquire a spindle-shape phenotype after 4 days.

**Fig. 1.**
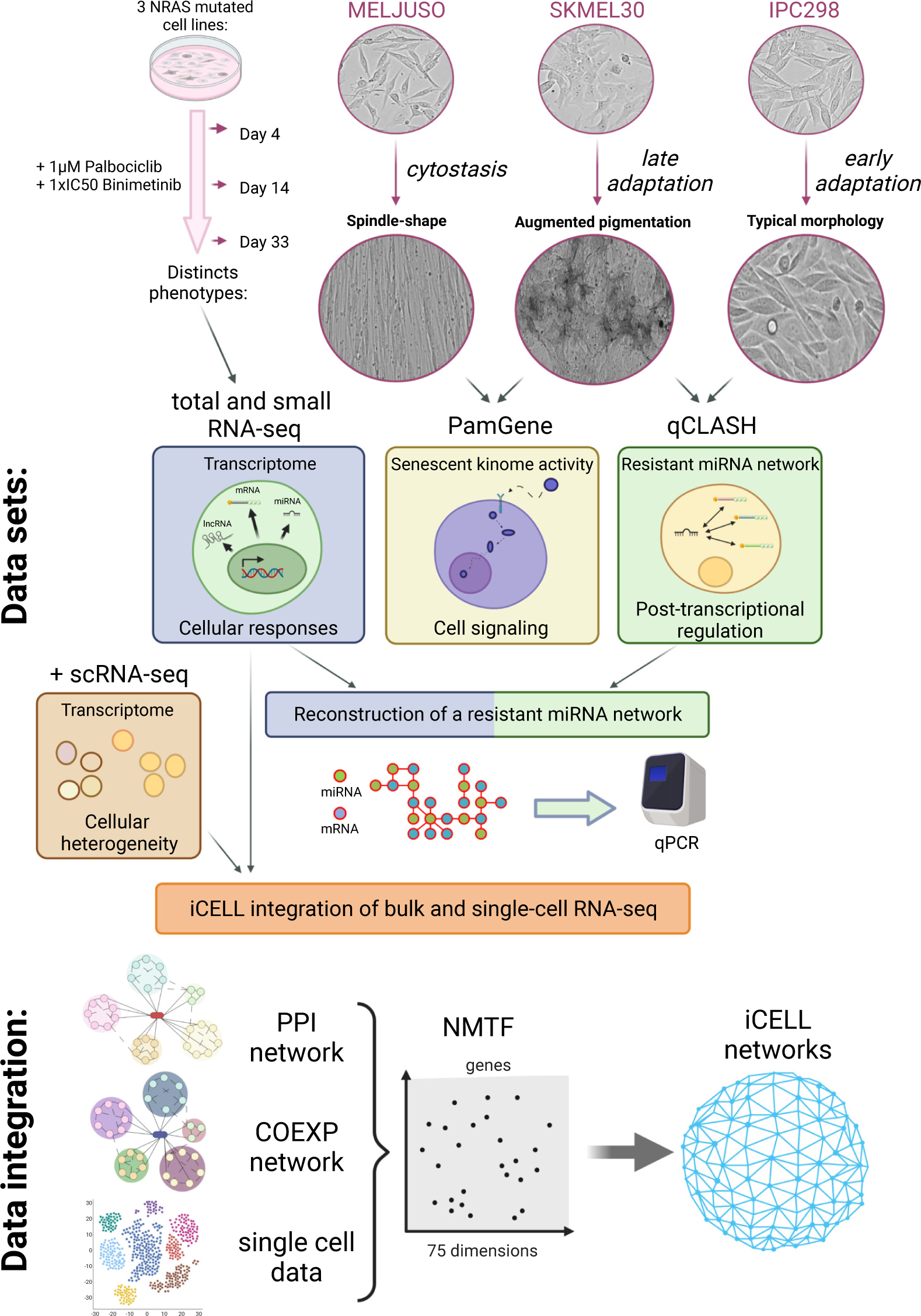
Experimental setup, cellular phenotypes and omics characterization. IPC298, MELJUSO and SKMEL30 cell lines display different phenotypes upon CDK4/6i and MEKi. We characterised those using RNA-seq, kinome profiling and qCLASH method. RNA-seq and qCLASH data were further combined to construct a resistant miRNA network. Bulk and single-cell RNA-seq were finally integrated into “iCell” networks.

As distinct morphological changes were observed at these time points, we profiled the transcriptome of the cell lines at day 0, 4, 14 and 33 using total RNA-seq. Cellular signaling supporting the adaptation to the treatment was investigated by profiling the kinome of the senescent MELJUSO and adaptative SKMEL30 cell lines using PamGene technology. Next, we characterised the resistant miRNA interactome of the SKMEL30 and IPC298 at 33 days by combining small RNA-seq and the qCLASH method. Finally, by integrating bulk and single cell RNA-seq data, deregulated pathways and predicted genes associated with senescence escape were identified.

### Cell lines display coherent transcriptomic profiles composed of interferons, EMT and senescent responses under prolonged CDK4/6i and MEKi treatment

Distinct cellular responses to treatment were reflected at the transcriptomic level where the cell lines clustered separately on the PCA (Fig. 2.A). As MELJUSO displayed a different evolution over time, we decided to analyze each cell line individually. Differential expression analysis, comparing later time points to day 0, revealed a greater change in the MELJUSO transcriptome. MELJUSO corresponds to 9548, 11362 and 10126 significant (p.adj <= 0.05) differentially expressed genes at day 4, day 14 and day 33, respectively when compared to SKMEL30 and IPC298 (9208, 9749, 6509 and 5436, 7575 and 3217, respectively) (Supp. Fig 1).

**Fig. 2.**
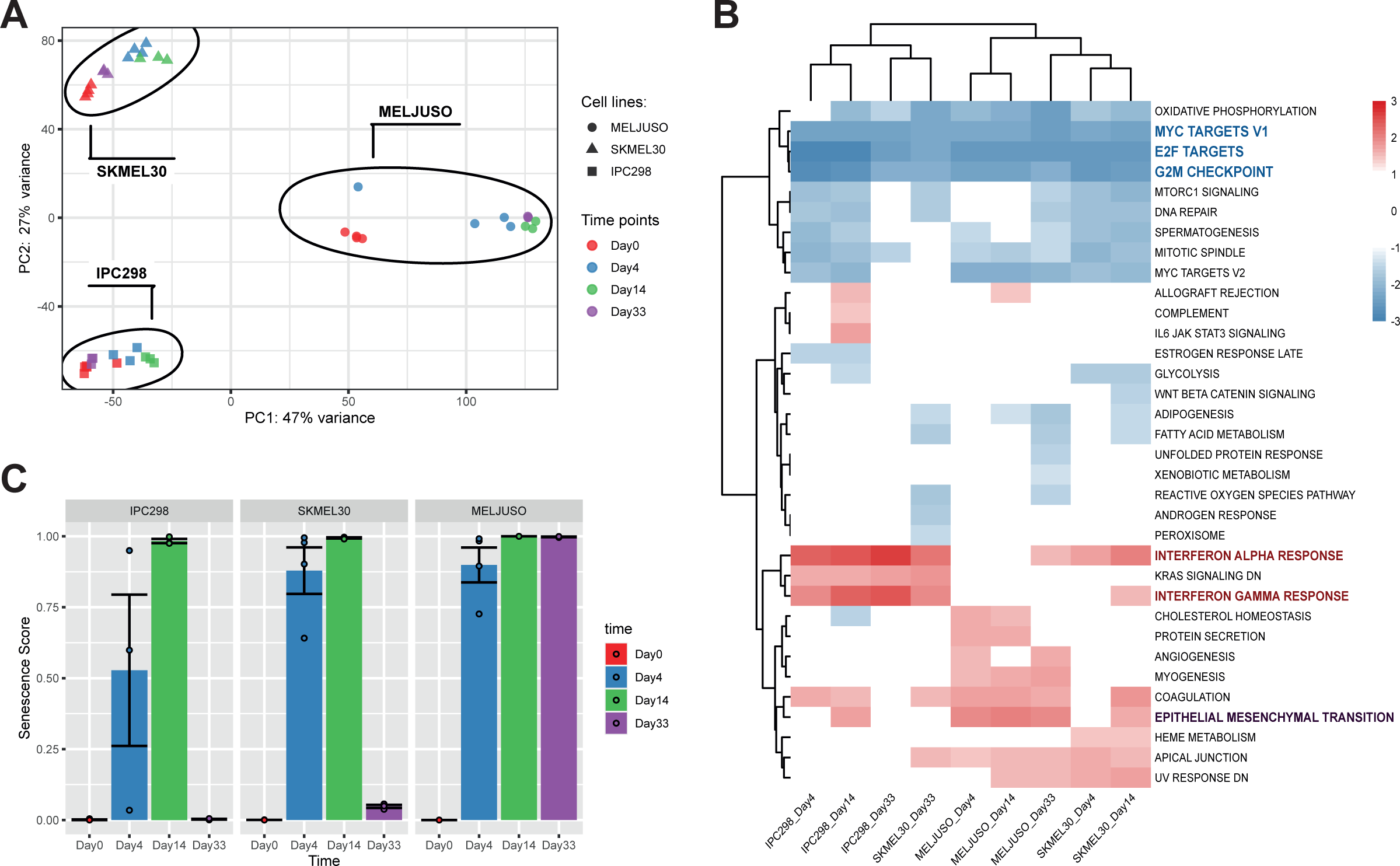
NRAS mutant melanoma trigger IFN and EMT responses upon MEKi and CDK4/6i. A. Principal component analysis clusters cell lines separately reflecting distinct observed phenotypes. B. Gene Set Enrichment Analysis using the hallmarks gene sets from MSigDB. All cell lines display an enrichment in interferons responses and at day 14 in EMT. C. Senescence scores predicted by Cancer SENESCopedia. SENESCopedia webtool (https://ccb.nki.nl/publications/cancer-senescence) uses a gene expression classifier to predict senescence in cancer cell samples [18].

Gene Set Enrichment Analysis (GSEA) was performed on differentially expressed genes using the “hallmarks” from MSigDB, which revealed the downregulation of pathways related to proliferation such as “MYC TARGETS V1”, “E2F TARGETS”, and “G2M CHECKPOINT” (Fig. 2.B), all affected by our inhibitors. The clustering of samples on top of the heatmap clearly separates proliferative samples (left part) comprising the IPC298 cell line and SKMEL30 day 33 from cytostatic samples (right part) with MELJUSO and early time points for SKMEL30. Interestingly, we noticed across all cell lines an enrichment in interferon responses and an enrichment in Epithelial Mesenchymal Transition (EMT) for the cytostatic samples (Fig 2.B). CDK4/6 inhibitors have been shown to induce cellular senescence in different cell types [16, 17]. We therefore used a recently published classifier, “SENESCopedia”, to predict the levels of senescence in our samples (Fig. 2.C) [18]. On day 14, all cell lines display high senescence scores suggesting that they all engage in a related transcriptional programme. Of note, all samples with a high senescence score are also enriched in EMT which is coherent with previous observations of EMT in different epithelial cell types after induction of senescence (fibroblasts, colorectal and lung cancer) [19, 20, 21].

Altogether, these results suggest that all 3 cell lines trigger the same type of responses composed of an interferon response, a senescence and “EMT” programme but with a different intensity and temporality, which may explain the different phenotypes observed.

### Kinome profiling of adapting cell lines shows reactivation of RTK pathways and suggests a key role of ERK5

To better understand how the cell lines adapt to the treatment, we acquired PamGene data to profile the kinome of SKMEL30 and MELJUSO cells at day 0, 1, 4 and 33 (Supp. Fig 2) as these two cell lines showed distinct responses to combined kinase inhibitors. MELJUSO stops proliferating and changes morphology after about 4 days while SKMEL30 acquires a dark pigmentation before restoring growth. Kinase activities were inferred from the levels of peptide phosphorylation at different time points and compared to day 0 (summarised in Fig 3.A). A phylogenetic representation shows that these two cell lines react similarly to the treatment only at early time points (Supp. Fig 3).

**Fig. 3.**
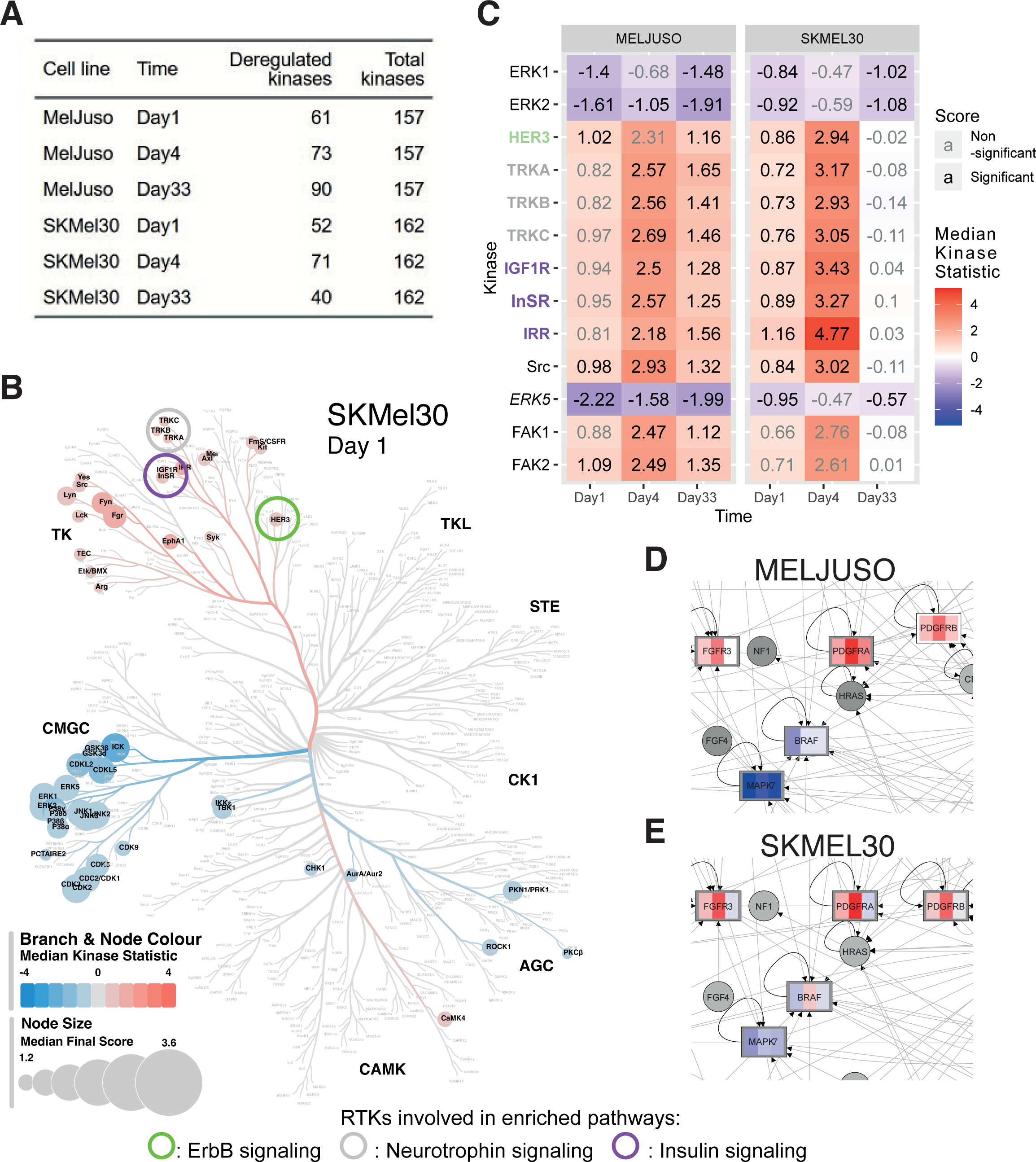
MEKi and CDK4/6i lead to RTKs reactivation. A. Summary of significantly deregulated kinases. B. Kinase activities after MEKi and CDK4/6i. ERK1/2 inhibition leads to RTKs activation. Rreceptors names are colored according to the enriched pathways. C. Phylogenetic tree for SKMEL30 on day 1. The size of the leafs represents the “Median Final Score”, a score greater than 1.2 indicates a significant change between conditions. The color of the branches and leaves shows the “Median Kinase Statistic”, which is the difference in kinase activity. Circles highlight RTKs contributing to enriched pathways. D. andE. Activity of ERK5 in the senescent and adaptative cell lines. D. zoom on the MAPK pathway showing the deregulation of ERK5 (*MAPK7*) in MELJUSO cell line. E. same for the SKMEL30 cell line.

To perform pathway enrichment on a reduced set of proteins such as kinases, we used a network-based approach and obtained coherent results for the two cell lines (Table 1). Indeed, pathways like RIG-I or Toll-like receptor signaling (related to innate immunity and interferon responses) were enriched at early time points for both SKMEL30 and MELJUSO. Among the enrichment results, RTK pathways involving ErbB, neurotrophin and insulin appear (Fig 3.B, 3.C). Some pathways were enriched before the increase in receptor activity reached the significance threshold and those were therefore not represented on the phylogenetic tree (Supp. Fig 3). This signaling seems to persist after the receptor resumes normal activity (Table 1). It has previously been shown that the inhibition of ERK1/2 suppresses negative feedback on RTK expression, which can then be further activated to compensate for this inhibition [22] (Fig 3 B, Supp. Fig 3). Coherent with the enriched pathways, we observed in both cell lines a significant increase in the activity of RTKs such as HER3 and NTRK2 as well as insulin-related receptors (INSR, IGF1R and IRR). Noteworthy, downstream insulin signaling was enriched in the cytostatic MELJUSO only (detailed representation of pathways Supp. Fig 4-8). Other RTKs previously related to resistance such ALK, AXL or c-MET displayed an increased activity (complete list Supp. Fig 9) but those were not associated with enriched downstream signaling. Within the MAPK pathway, the decrease in ERK5 (*MAPK7*) activity was partially relieved in SKMEL30 but not in MELJUSO (Fig 3.B). Along these lines, Tubita et al recently showed that knock-down by shRNA or inhibition of the kinase activity of ERK5 triggers senescence in melanoma [23].

**Table 1.** Enrichment results from PamGene data. Network-based enrichment was performed on the significantly deregulated kinases at different time points. MELJUSO and SKMEL30 cells show at all time points an enrichment in ErbB and neurotrophin pathway. Only MELJUSO cells show an enrichment in insulin pathway at day 4 and day 33.

| Pathway enrichment results for MELJUSO cell line |  |  |  |  |  |  |  |  |
| --- | --- | --- | --- | --- | --- | --- | --- | --- |
| Day1 |  |  | Day4 |  |  | Day33 |  |  |
| Pathway Name | XD-score | p-adj | Pathway Name | XD-score | p-adj | Pathway Name | XD-score | p-adj |
| Shigellosis | 1.83 | 6.12E-20 | mTOR signaling pathway | 1.13 | 1.33E-10 | Type II diabetes mellitus | 1.42 | 1.49E-09 |
| NOD-like receptor signaling pathway | 1.38 | 6.69E-14 | Acute myeloid leukemia | 1.05 | 1.50E-09 | mTOR signaling pathway | 1.08 | 2.57E-11 |
| Epithelial cell signaling in Helicobacter pylori infection | 1.12 | 3.49E-13 | Type II diabetes mellitus | 0.89 | 3.91E-06 | **ErbB signaling pathway** | 1.01 | 3.31E-12 |
| Progesterone-mediated oocyte maturation | 1.02 | 1.41E-14 | Glioma | 0.86 | 8.12E-08 | Acute myeloid leukemia | 1.00 | 3.13E-10 |
| RIG-I-like receptor signaling pathway | 1.00 | 1.99E-11 | **ErbB signaling pathway** | 0.86 | 2.02E-10 | Fc epsilon RI signaling pathway | 0.95 | 4.51E-12 |
| **ErbB signaling pathway** | 0.97 | 2.54E-13 | Gap junction | 0.77 | 3.34E-09 | Progesterone-mediated oocyte maturation | 0.95 | 4.23E-12 |
| GnRH signaling pathway | 0.93 | 1.47E-14 | Prion diseases | 0.75 | 2.78E-04 | Glioma | 0.95 | 1.69E-08 |
| Type II diabetes mellitus | 0.91 | 8.02E-07 | **Adherens junction** | 0.72 | 1.69E-07 | Prion diseases | 0.95 | 2.91E-05 |
| Fc epsilon RI signaling pathway | 0.90 | 6.70E-13 | Fc gamma R-mediated phagocytosis | 0.71 | 2.99E-10 | Adipocytokine signaling pathway | 0.87 | 2.60E-08 |
| **Neurotrophin signaling pathway** | 0.89 | 4.52E-17 | **VEGF signaling pathway** | 0.68 | 5.13E-08 | **VEGF signaling pathway** | 0.86 | 5.62E-10 |
| Toll-like receptor signaling pathway | 0.89 | 5.76E-14 | Fc epsilon RI signaling pathway | 0.65 | 7.58E-09 | Shigellosis | 0.86 | 1.53E-07 |
| Pancreatic cancer | 0.76 | 1.12E-08 | Prostate cancer | 0.65 | 5.45E-08 | **Neurotrophin signaling pathway** | 0.83 | 5.38E-13 |
| Colorectal cancer | 0.75 | 1.52E-07 | Long-term potentiation | 0.65 | 1.48E-06 | Aldosterone-regulated sodium reabsorption | 0.82 | 4.81E-05 |
| T cell receptor signaling pathway | 0.73 | 2.23E-12 | Aldosterone-regulated sodium reabsorption | 0.64 | 3.96E-04 | Gap junction | 0.82 | 7.28E-11 |
| **VEGF signaling pathway** | 0.69 | 6.13E-09 | Tight junction | 0.62 | 6.96E-11 | Prostate cancer | 0.80 | 1.21E-09 |
| Acute myeloid leukemia | 0.57 | 9.12E-08 | Vascular smooth muscle contraction | 0.60 | 4.24E-09 | **Adherens junction** | 0.79 | 3.65E-08 |
| Prostate cancer | 0.56 | 9.34E-08 | **Insulin signaling pathway** | 0.59 | 1.38E-10 | GnRH signaling pathway | 0.77 | 9.13E-11 |
| Leishmaniasis | 0.48 | 6.22E-06 | **Neurotrophin signaling pathway** | 0.59 | 2.88E-10 | T cell receptor signaling pathway | 0.75 | 3.13E-11 |
| Dorso-ventral axis formation | 0.48 | 1.13E-02 | Progesterone-mediated oocyte maturation | 0.58 | 8.17E-07 | Long-term potentiation | 0.72 | 2.77E-07 |
|  |  |  | GnRH signaling pathway | 0.55 | 6.47E-08 | NOD-like receptor signaling pathway | 0.71 | 1.45E-06 |
|  |  |  | Adipocytokine signaling pathway | 0.53 | 2.89E-05 | Pancreatic cancer | 0.69 | 2.77E-07 |
|  |  |  | Hedgehog signaling pathway | 0.52 | 7.95E-05 | Fc gamma R-mediated phagocytosis | 0.65 | 1.68E-09 |
|  |  |  | Long-term depression | 0.51 | 7.43E-07 | Toll-like receptor signaling pathway | 0.64 | 4.58E-09 |
|  |  |  | T cell receptor signaling pathway | 0.47 | 2.18E-07 | Epithelial cell signaling in Helicobacter pylori infection | 0.63 | 2.77E-07 |
|  |  |  | Chemokine signaling pathway | 0.44 | 4.38E-14 | Long-term depression | 0.59 | 1.26E-07 |
|  |  |  |  |  |  | Vascular smooth muscle contraction | 0.55 | 2.33E-08 |
|  |  |  |  |  |  | **Insulin signaling pathway** | 0.54 | 9.87E-10 |
|  |  |  |  |  |  | Colorectal cancer | 0.54 | 3.31E-05 |

RTKs involved in enriched pathways: ○ : ErbB signaling ○ : Neurotrophin signaling ○ : Insulin signaling
| Pathway enrichment results for SKMEL30 cell line |  |  |  |  |  |  |  |  |
| --- | --- | --- | --- | --- | --- | --- | --- | --- |
| Day1 |  |  | Day4 |  |  | Day33 |  |  |
| Pathway Name | XD-score | p-adj | Pathway Name | XD-score | p-adj | Pathway Name | XD-score | p-adj |
| Shigellosis | 1.27 | 1.09E-13 | Acute myeloid leukemia | 1.27 | 8.19E-10 | Shigellosis | 1.17 | 1.64E-11 |
| Fc epsilon RI signaling pathway | 1.17 | 2.92E-16 | Glioma | 0.91 | 4.31E-08 | NOD-like receptor signaling pathway | 1.02 | 3.88E-10 |
| NOD-like receptor signaling pathway | 1.12 | 5.18E-11 | mTOR signaling pathway | 0.83 | 1.43E-09 | Dorso-ventral axis formation | 0.94 | 2.69E-04 |
| Progesterone-mediated oocyte maturation | 0.95 | 1.99E-13 | **ErbB signaling pathway** | 0.80 | 1.52E-09 | Pancreatic cancer | 0.85 | 1.06E-09 |
| Type II diabetes mellitus | 0.94 | 4.51E-07 | Type II diabetes mellitus | 0.74 | 4.02E-05 | Type II diabetes mellitus | 0.80 | 2.89E-06 |
| RIG-I-like receptor signaling pathway | 0.91 | 3.10E-10 | Adipocytokine signaling pathway | 0.71 | 1.06E-06 | Epithelial cell signaling in Helicobacter pylori infection | 0.79 | 1.70E-08 |
| Epithelial cell signaling in Helicobacter pylori infection | 0.88 | 2.11E-10 | Long-term potentiation | 0.69 | 7.97E-07 | Progesterone-mediated oocyte maturation | 0.78 | 4.11E-10 |
| GnRH signaling pathway | 0.87 | 2.04E-13 | Fc gamma R-mediated phagocytosis | 0.66 | 2.29E-09 | Fc epsilon RI signaling pathway | 0.75 | 1.53E-09 |
| **Neurotrophin signaling pathway** | 0.86 | 3.07E-16 | Gap junction | 0.60 | 3.74E-07 | GnRH signaling pathway | 0.74 | 1.54E-11 |
| **ErbB signaling pathway** | 0.80 | 9.43E-11 | Prostate cancer | 0.59 | 3.74E-07 | **ErbB signaling pathway** | 0.65 | 5.44E-09 |
| Toll-like receptor signaling pathway | 0.75 | 1.70E-11 | Fc epsilon RI signaling pathway | 0.58 | 6.98E-08 | Toll-like receptor signaling pathway | 0.62 | 1.04E-09 |
| **VEGF signaling pathway** | 0.73 | 2.85E-09 | Vascular smooth muscle contraction | 0.57 | 2.42E-08 | RIG-I-like receptor signaling pathway | 0.58 | 6.16E-07 |
| Aldosterone-regulated sodium reabsorption | 0.69 | 9.10E-05 | Prion diseases | 0.54 | 2.70E-03 | T cell receptor signaling pathway | 0.57 | 1.31E-09 |
| T cell receptor signaling pathway | 0.69 | 2.45E-11 | **Adherens junction** | 0.52 | 1.93E-05 | Leishmaniasis | 0.57 | 6.62E-07 |
| Leishmaniasis | 0.65 | 1.91E-07 | **Neurotrophin signaling pathway** | 0.49 | 2.15E-08 | Colorectal cancer | 0.54 | 9.30E-06 |
| **Adherens junction** | 0.65 | 2.25E-07 | Apoptosis | 0.49 | 4.20E-06 | Endometrial cancer | 0.51 | 9.97E-05 |
| Colorectal cancer | 0.64 | 1.78E-06 | Tight junction | 0.46 | 5.47E-08 | **Neurotrophin signaling pathway** | 0.47 | 3.13E-09 |
| Prion diseases | 0.54 | 1.08E-03 | Non-small cell lung cancer | 0.46 | 7.94E-05 | **VEGF signaling pathway** | 0.42 | 8.77E-06 |
| Dorso-ventral axis formation | 0.52 | 8.88E-03 | Vibrio cholerae infection | 0.46 | 7.74E-04 |  |  |  |
| Amyotrophic lateral sclerosis (ALS) | 0.48 | 2.12E-04 | Aldosterone-regulated sodium reabsorption | 0.46 | 3.50E-03 |  |  |  |
|  |  |  | Long-term depression | 0.43 | 6.55E-06 |  |  |  |
|  |  |  | Salivary secretion | 0.42 | 6.05E-05 |  |  |  |
|  |  |  | Progesterone-mediated oocyte maturation | 0.41 | 6.05E-05 |  |  |  |
|  |  |  | Gastric acid secretion | 0.40 | 2.79E-04 |  |  |  |
|  |  |  | **VEGF signaling pathway** | 0.37 | 1.14E-04 |  |  |  |

Overall, the kinome data show the reactivation of ErbB, neurotrophin and insulin pathways and enrichment of their downstream signaling. Interestingly, ERK5 activity was strongly reduced in the cytostatic cell line MELJUSO (Fig 3.B).

### Contribution of miRNAs to the resistant phenotype

To see if miRNAs could participate in the adaptation to treatment and the proliferative phenotypes, we profiled the miRNome of all three cell lines by small RNA-seq at day 0 and 33. In line with their phenotypes, IPC298 showed little deregulation of miRNAs compared to SKMEL30 and MELJUSO cell lines (Supp Fig. 10). Next, we performed qCLASH analysis for the two proliferative cell lines IPC298 (early adaptation) and SKMEL30 (late adaptation) (summarised in Fig 4.A).

**Fig. 4.**
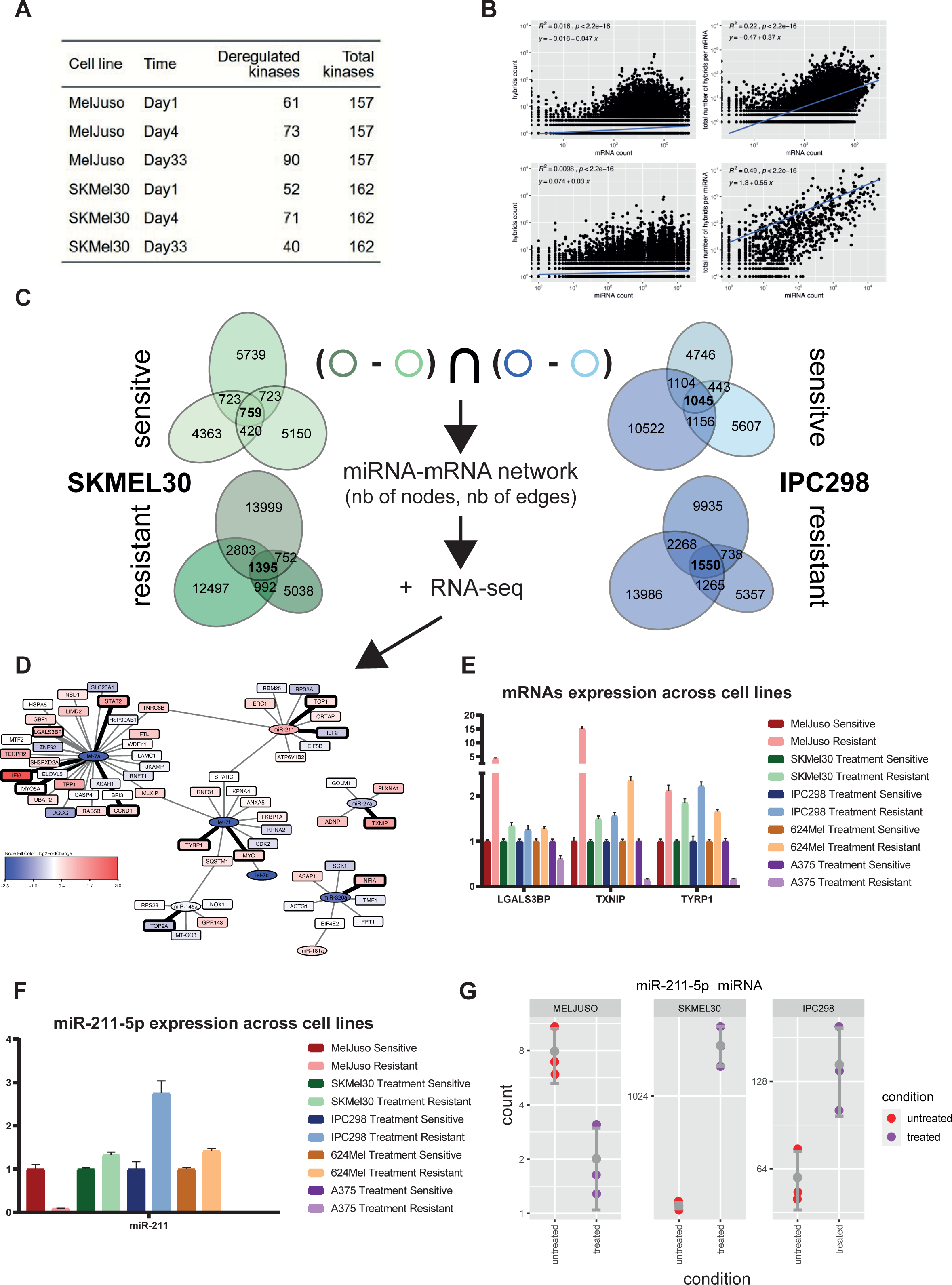
miRNA-mRNA interactions potentially contributing to resistance. A. Summary of the interactions detected through the qCLASH method. B. Top part, correlations between hybrid counts (qCLASH) and miRNA/mRNA counts (RNA-seq). Bottom part, summing hybrid counts per mRNA or miRNA (qCLASH) further strengthen the correlations with the RNA-seq. To select candidates, we considered the interactions detected in three technical replicates (colored circles). Then for the two cell lines, interactions present in the resistant state but absent in the sensitive state were considered. Finally, interactionscommon to the two cell lines IPC298 and SKMEL30 were used. D. RNA-seq data for SKMEL30 cell line as log fold changes overlayed onto the resistant miRNA network. Selected interactions for further validation by qPCR are represented with thick lines and the corresponding mRNAs and miRNAs with bolded borders. E. Relative expression of LGALS3BP, TXNIP and TYRP1 mRNAs assessed by qPCR across three NRAS and two BRAF mutant melanoma cell lines. F. Relative expression of miR-211-5p across the same cell lines.

Although qCLASH-based miRNA interactomes do not provide a complete picture of all interactions, PCA was able to discriminate between the different cell lines and conditions (Supp. Fig 11). Previous studies reported a correlation between the number of detected hybrids and the expression levels of its miRNA and mRNA components [14, 24]. These correlations hold true for our matching total and the small RNA-seq data when taking the total count across the replicates (Fig 4.B, left panels). Moreover, further summing up the hybrids for each miRNA or mRNA strengthens the correlation between RNA-seq and qCLASH data, especially for miRNAs (Fig. 4. B, right panels). qCLASH data also contain transient interactions, which are unlikely to be functionally relevant for the cell and technical replicates are often used to further select most likely interactions. We explored how the detection of interactions in one, two or three replicates depends on their expression level and observed that interactions detected in triplicates were associated with a greater expression supporting the rationale of using such thresholds (Supp. Fig 12). To further select the most relevant miRNAs, we constructed a network with the interactions specific to the resistant phenotype. We first selected the interactions detected in triplicates, then ensured that they were present in both resistant IPC298 and SKMEL30 but absent from their untreated counterparts (Fig 4.C). Finally, we overlaid the log fold changes from the small and total RNA-seq for the corresponding miRNAs and mRNAs onto the resulting network (Fig. 4.D). Based on those representations, we selected some interactions to confirm their deregulation by qPCR. We investigated if those observations could be extended to two BRAF mutant cell lines, A375 and 624Mel which underwent the same treatment and re-established growth (Supp. Fig 13). Most of the selected interactions did not exert a robust inverse expression between miRNA and mRNA (Supp. Fig 14,15). Among mRNAs, we observed a robust upregulation of CCND1 which is a known resistance mechanism to CDK4/6i as well as a strong upregulation of IFN related gene IFI6 (Supp. Fig 13). Except for the amelanotic cell line A375, we observed the upregulation of LGALS3BP, TXNIP and TYRP1 (Fig 4.E). Among miRNAs, mir-211-5p was upregulated in all proliferative cell lines except the amelanotic A375, which is known not to express it (Fig 4.F) [25], and MELJUSO (Fig 4.G), which has very low levels.

Overall, the number of interactions detected by qCLASH method is influenced by the expression level of the corresponding miRNA and mRNA. Most of the detected interactions did not exert the expected inverse expression pattern between miRNA and mRNA. miRNAs and mRNAs displayed a more robust change in expression across cell lines. Except for the amelanotic A375, upregulation of miR-211-5p is associated with resistant and proliferative cell lines.

### iCell integration of bulk and single-cell RNA-seq reveals perturbed pathways and predicts genes facilitating senescence escape

To gain further insights into NRAS mutant melanoma cells’ adaptation to treatment, we integrated the bulk RNA-sequencing data with previous condition-matching single-cell data from our lab by adapting and applying a Non-negative Matrix Tri-Factorization (NMTF) approach called iCell [15]. We used the iCell methodology because it is a versatile data fusion framework, which has previously been used to perform integrative comparative analyses of disease and control tissue data, predicting new cancer and COVID-19 related genes [15, 26] and suggesting potential drug re-purposing candidates [26].

For each condition (treated cell line and time point), we constructed protein-protein interaction (PPI) networks by overlaying the experimentally validated PPI network for *Homo Sapiens* from BioGRID [27] with the condition-specific bulk RNA-sequencing data. Also, we derived a condition-specific co-expression (COEXP) network from the corresponding bulk RNA-seq data (see Methods section). Next, we applied Non-negative Matrix Tri-Factorization (NMTF) on the corresponding adjacency matrices of the resulting PPI, COEXP and the single-cell expression data collectively, to ensure data fusion (Methods and [15]). This resulted in the creation of nine condition-specific iCell networks, by multiplying the G1 matrix resulting from the data fusion and its transpose (which is equivalent to computing the dot product between gene pairs) and considering as interactions the 1% highest values in each row and column (see the Methods and [15]).

As illustrated in Supp. Fig 16, resulting condition-specific iCell networks were investigated by first comparing the global topology of all resulting iCell networks over all treated single-cell line conditions over all time points. Then, these topological changes were further characterised by identifying the genes that change their local topology between each pair of conditions, as well as the ones whose local topology is conserved. We related the identified most rewired genes and the least rewired ones to MSigDB hallmarks by performing an Over-Representation Analysis (ORA) (see Methods for details). For each condition, we also clustered the genes based on the local topology in the corresponding condition-specific iCell network and then found the MSigDB hallmarks that are enriched in the resulting clusters. We focused on two senescence-escape related hallmark pathways to predict novel genes potentially related to these pathways based on their appearance in the clusters that were enriched across all conditions.

Then, we compared the overall topology of the iCell networks obtained for each condition by using thus far the most sensitive measure of global network topology, the “Graphlet Correlation Distance” (GCD-11) (see Methods and [28]). The lower the GCD-11 value, the closer the topologies of the networks [28]. We used Ward’s hierarchical clustering to cluster the resulting heatmap of the GCD-11s between all iCell networks (all-to-all matrix). In line with our phenotypic observations, we showed on the heatmap (Fig 5.A) that for the resistant cell line IPC298, the conditions D4 and D33 have the lowest GCD-11 value (0.26) and hence clustered together, suggesting early adaptation to treatment of this cell line. For the two other cell lines, conditions D0 and D4 grouped together (GCD-11 values of 0.71 for SKMEL30 and 0.87 for MELJUSO), suggesting that these two cell lines adapt later.

**Fig. 5.**
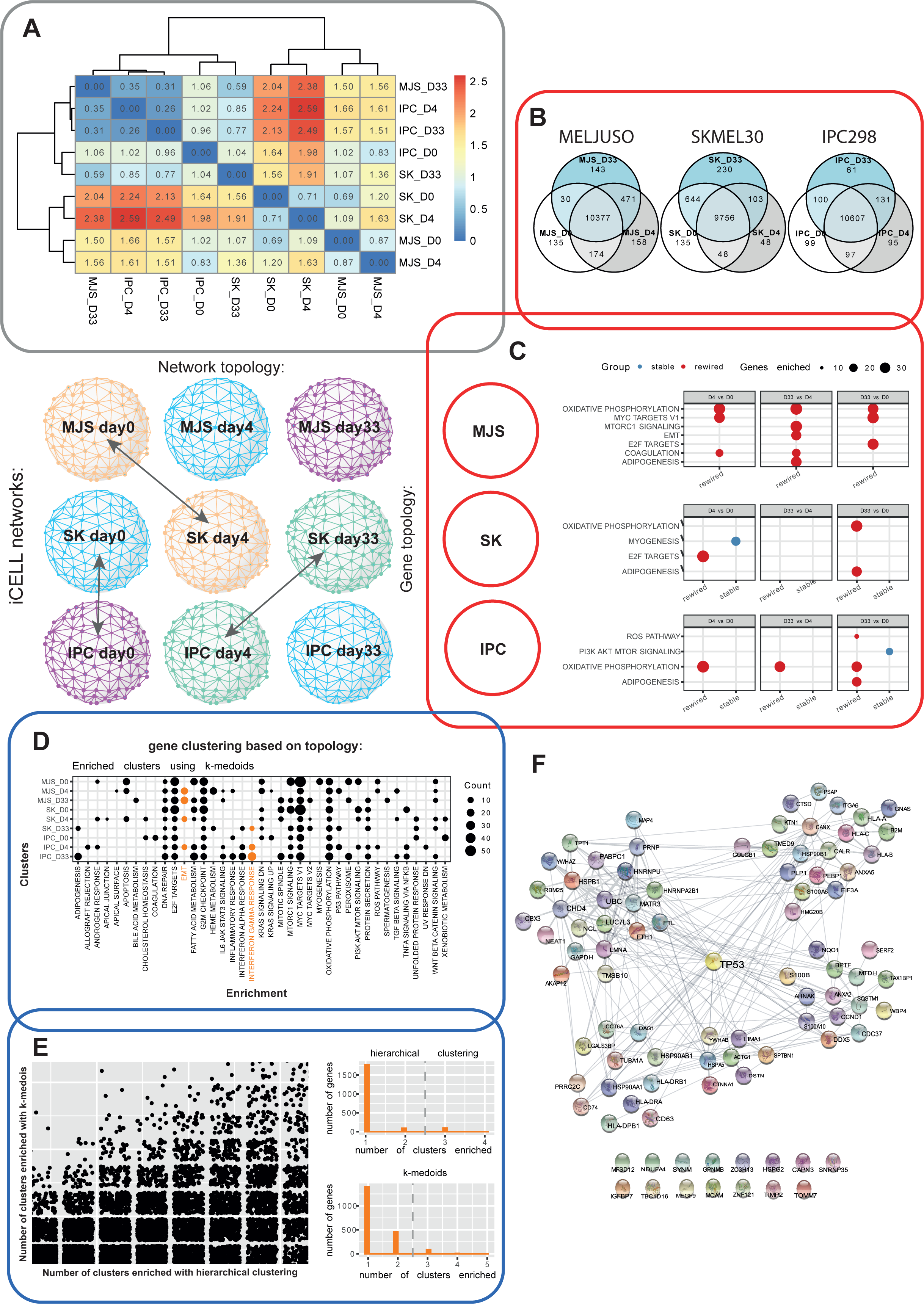
“iCell” integration of bulk and single-cell RNA-seq. A. Comparison of iCell networks topology using Graphlet Correlation Distances (GCD). iCELL networks recapitulate phenotypic observations and early adaptation of IPC298. B. Venn diagrams representing genes overlap between conditions. C. Comparison of gene local topology using Graphlet Degree Signature Similarity (GDSS). Within all cell lines, the different time points were compared. ORA was performed on the top 10% most “stable” or “perturbated” genes. D. Genes were clustered according to their Graphlets Degree Signatures. ORA was performed to associate gene clusters with biological function. E. Left panel: number of enriched clusters detected for each gene across all conditions. Right panel, Number of detected clusters for the EMT and iFN gamma pathways (related to senescence escape) for the two algorithms. F. Protein-protein interactions (BioGRID) between P53 and the genes associated with EMT and IFN gamma.

Next, the local topology of genes in the condition-specific iCell networks was analysed to find out if some are rewired under specific biological conditions. To this end, when comparing two conditions, we focused on genes that were expressed in both conditions (Venn diagrams in Fig. 5.B and supp. Fig. 17). Within each iCell network, the local topology around a gene in the network was quantified by using it’s “Graphlet Degree Vector” (GDV). We then measure the rewiring of a gene between two conditions by comparing its GDVs in the two networks using “Graphlet Degree Vector Similarity” (GDVS) (detailed in Methods and in [29]). Between two conditions, genes with high GDVSs have conserved topologies and are called “stable”, while the genes with low GDVSs are rewired and are called “perturbed”. We related the top 10% of the most “stable” and the top 10% of the most “perturbed” genes to MSigDB hallmarks [30] by performing ORA. For each cell line, the condition-specific iCell networks were compared for all time points together (i.e., D4 vs D0, D33 vs D4, and D33 vs D0). For all cell lines, we observed that the top 10% of the most “perturbed” genes were enriched at different contrasts in “ADIPOGENESIS” (also observed in senescent fibroblasts [31]) and “OXIDATIVE PHOSPHORYLATION” (OxPhos) (a pathway commonly dysregulated in senescence which has been proposed as a targeting strategy [32]) (Fig 5.C). Then, the local topology of genes was compared between the condition-specific iCell networks of different cell lines, but at matching time points (e.g., MELJUSO_D0 vs SKMEL30_D0). Interestingly, the top 10% of the most “stable” genes were only found to be enriched when comparing IPC298 vs MELJUSO at day 0 and when comparing MELJUSO vs SKMEL30 at day 4, which suggests that these cell lines react coherently early on and diverge later (supp Fig. 17). At day 0, and across all pairwise comparisons involving IPC298 cell line, the top 10% of the most “perturbed” genes were enriched for OxPhos, suggesting a different initial metabolic state for IPC298. On day 4, when comparing IPC298 and SKMEL30 cells, the top 10% of the most “perturbed” genes were enriched in “INTERFERON ALPHA/GAMMA RESPONSE”. At days 4 and 33, but not across all pairwise comparisons, we found that the top 10% of the most “perturbed” genes are enriched in “EPITHELIAL MESENCHYMAL TRANSITION”, “ADIPOGENESIS” and “COAGULATION”. Further information on the relevance of these enriched pathways can be found in the Discussion.

Finally, to predict novel genes which could contribute to senescence escape facilitated by interferons, genes were clustered based on their GDVs signatures in the corresponding condition-specific iCell network using “Ward’s hierarchical clustering” and the “k-medoids” algorithms. For each condition, we performed ORA to associate the clusters to hallmarks from MSigDB (results of ORA on the clusters obtained by using k-medoids clustering in Fig 5.D). The results for the two algorithms showed limited agreement (Fig 5.E, left), as measured by Adjusted Rand Index for each condition: 0.31, 0.38 and 0.30 for MELJUSO at day 0, day 4 and day33; 0.32, 0.30 and 0.26 for SKMEL30 at the same time points and 0.27, 0.31 and 0.45 for IPC298. Therefore, we considered the results from the two different clustering methods (hierarchical and k-medoids) separately. Enriched clusters were filtered to exclude genes that are already known to be associated with hallmarks in MSigDB. For the remaining genes, we counted the number of times a gene was found in the clusters that are enriched in one of the two senescence escape-related pathways, EMT and interferon gamma (see Fig 5.E right). Then, based on the distributions of the number of times genes appeared in these enriched clusters, we predicted the genes that appeared in two or more of these enriched clusters as related to senescence escape. To obtain more robust predictions, we only considered genes to be involved senescence escape-related by using both algorithms. This resulted in a set of 90 genes, some of which are known to interact with TP53, further suggesting a connection to senescence escape (Fig. 5.F, Supp. Table 2).

To conclude, the integrated analysis of bulk and single-cell data by using the iCell methodology further confirms our phenotypic observations and uncovers the most dysregulated processes (“OXIDATIVE PHOSPHORYLATION”, “ADIPOGENESIS”, “COAGULATION” and “EPITHELIAL MESENCHYMAL TRANSITION”) during adaptation to the treatment. Furthermore, it allowed us to predict 90 new genes involved in senescence escape.

## DISCUSSION

Most cancer cells inevitably develop resistance to targeted therapies. Anticipating the development of such resistance to a newly tested combination of CDK4/6i and MEKi, we characterised the transcriptomic responses in 3 NRAS mutant melanoma cell lines over an extended treatment duration of 33 days. Our RNA-seq data exhibited a strong enrichment in interferon responses, coherent with previous observations by Goel et al. [33] following CDK4/6 inhibition. This study showed that CDK4/6 inhibitors downregulate E2F2 and DNMT1, subsequently leading to a hypomethylation of the genome (Supp. Fig 18). The works of Chiappinelli et al [34] and Roulois et al [35] first showed that hypomethylation brought about by DNMT inhibitors induce the expression of otherwise silenced endogenous retroviruses (ERVs). These ERVs then form dsRNAs in the cytoplasm and trigger an anti-viral immune response associated with interferon production. This phenomenon was termed “viral mimicry” and has in the meantime been reported in different cancer types and in response to different epigenetic perturbations (DNMTi, HDACi, HMTi). Brägelmann et al [36] also reported the activation of viral mimicry in the case of BRAF mutant melanoma and MAPK inhibitors (BRAFi + MEKi) further supporting the E2F2-DNMT1 axis as a mechanism of action. Melanoma cells are considered very immunogenic [37] and this partially explains why melanoma respond better to immunotherapy than other cancer types. IFN signaling is known to upregulate PD-L1, explaining why many treatments activating viral mimicry have been shown to synergise with immune checkpoint inhibitors. Here, we confirm the appearance of an IFN signature following CDK4/6 and MEK inhibition suggesting a potential induction of viral mimicry.

A “senescence score” predicted from our data with SENESCopedia [18] suggested induction of a senescent transcriptional programme in all cell lines. Cytokines composing the SASP can induce features of EMT [38] and EMT has been recently proposed to inhibit senescence in HeLa cells treated with Doxorubicin [39]. TIS and EMT share common transcription factors with opposite effects on both processes (Supp Fig. 19) [40] and cancer cells escaping from TIS have been associated with a mesenchymal phenotype [41]. IFN gamma, by promoting EMT [42, 43, 44], could further support escape from senescence (Supp Fig. 20.A). To understand the relationship between IFN gamma and EMT gene sets (from the hallmarks) and a recent gene set characterizing senescence in cancer cell lines [45], we looked at changes in expression (log2 fold change) between those gene sets at day 14 where all cell lines display high senescence scores. IFN gamma shows an opposite trend in expression to the EMT and senescence gene sets (Supp Fig. 20.B), suggesting that IFN may accelerate the EMT process and thereby repress senescence. Albeit interferon signaling has earlier on been linked with senescence [45], only later Katlinskaya and Yu et al. showed a role of IFNAR1 in the induction of senescence [47]. Recently, the accumulation of double stranded RNA was proposed to also contribute to the onset of senescence [48, 49]. These findings suggest a link between senescence and viral mimicry, which can engage autocrine and paracrine secretion of interferons. We propose that the induced IFN gamma programme may contribute to senescence escape by inducing EMT.

To identify targetable proteins, which could explain the senescent phenotype, we profiled the kinase activities of SKMEL30 and MELJUSO cells upon CDK4/6 and MEK inhibition. We observed the reactivation of different RTKs due to ERK1/2 inhibition as previously reported by others [50]. Some of the corresponding pathways such as ErbB, neurotrophin and insulin were found enriched through network-based enrichment hinting at an active downstream signalling. Interestingly, all those pathways have been shown to activate ERK5. In breast cancer, ERK5 contributes to the response to EGF stimulation [51] and is involved in the resistance to ErbB-2 (HER2) inhibitors by facilitating G1-S transition [52]. In BRAF mutant melanoma resistant to BRAFi and MEKi, IGF1R activation induces ERK5 phosphorylation [53]. In senescent fibroblasts, BDNF-TrkB has been shown to activate ERK5 and was proposed to form an autocrine loop further enhancing senescent cell viability [54]. ERK5 activation has also been linked to PDGFR signalling in NRAS mutant melanoma treated with MEK and ERK1/2 inhibitors [55, 56] (Fig 3B, 3D, 3E). In other cell types, Src is also known to activate ERK5 and Focal Adhesion Kinases (FAK1 and FAK2) [57, 58]. Other Src Family Kinases (SFKs) display an increased activity (Supp. Fig 20) and have been previously linked with EMT [59]. Finally, IFN gamma was reported to activate ERK5 and its activation is necessary for the full expression of interferon stimulated genes [60]. Linking this observation with our RNA-seq data (Supp. Fig. 21) further supports the idea that IFN gamma could facilitate senescence escape by activating ERK5.

Investigating the contribution of miRNAs to the establishment of resistance, we noticed the upregulation of mir-211-5p in all proliferative cell lines but the amelanotic A375. Interestingly, miR-211-5p is known to derive from the TRPM1 transcript and both are very lowly expressed in the senescent cell line MELJUSO (supp Fig. 21). miR-211-5p inhibits DUSP6, which regulates ERK5 kinase activity, and this axis has been shown to be relevant i*n vivo* in genetic variants of A375 overexpressing mir-211 or DUSP6 and treated with BRAF/MEKi [61]. In qPCR validation experiments, mRNAs displayed a more robust change in expression with LGALS3BP, TXNIP and TYRP1 being upregulated in four out of five cell lines. LGALS3BP and TXNIP are interferon-inducible and interestingly, LGALS3BP has been reported to promote EMT.

Integrating single-cell and bulk RNA-seq data, we observed the perturbation of OxPhos in all cell lines. This is in line with previous works demonstrating metabolic adaptation after MAPK inhibition or CDK4/6i in BRAF mutant and uveal melanoma [62, 63]. Moreover, OxPhos metabolism has been previously reported as a generic drug resistance mechanism and as a key decision point between cell death and survival [64, 65]. Upon comparison of different treatment time points, differences in processes such as “ADIPOGENESIS” and “COAGULATION” were observed. Interestingly, those have recently been associated with replicative senescence [38], cell cycle arrest caused by CDK4/6i [66] and escape from OIS [67]. EMT has been reported to inhibit senescence [22, 44]. We therefore clustered the genes in the networks and enriched those clusters to report novel associations between genes and the processes of EMT and IFN gamma response, which could support senescence escape. Our predicted genes are coherent with the results of a genetic screen aiming to identify genes contributing to senescence bypass after p16 overexpression in melanoma [68]. Interestingly, the IFN-inducible gene LGALS3BP, previously reported to contribute to EMT and identified in our miRNA resistant network, was predicted by our approach to be associated with senescence escape. On the other hand, TXNIP, another IFN-inducible gene from the miRNA network, recently shown to participate in the induction of senescence upon IGF1 stimulation, was found associated only once and therefore not selected [69]. This further suggests that our approach can relate genes to function beyond their expression pattern, providing a list of 90 genes associated with senescence escape in cancer cells (Fig. 5.F).

Taken together, our data describe how NRAS mutant melanoma adapt to CDK4/6 and MEK inhibition by triggering an EMT programme. This is reminiscent of the Neural Crest Stem Cell (NCSC)-like state identified in single cell RNA-seq of patient samples after MAPKi by Rambow and colleagues [70]. This state was later characterised by FAK-dependent AKT activation [71], associated with slow or no proliferation and the expression of EMT-related transcription factors.

In sum, our work sheds new light on the intimate relationships between three interacting pathways, senescence, interferon and insulin signalling, with the EMT process (Fig. 6). Our data suggest a new role of interferon gamma, which, by triggering EMT, can facilitate senescence escape while insulin signaling is associated with a robust cytostasis and senescent transcriptome.

**Fig. 6.**
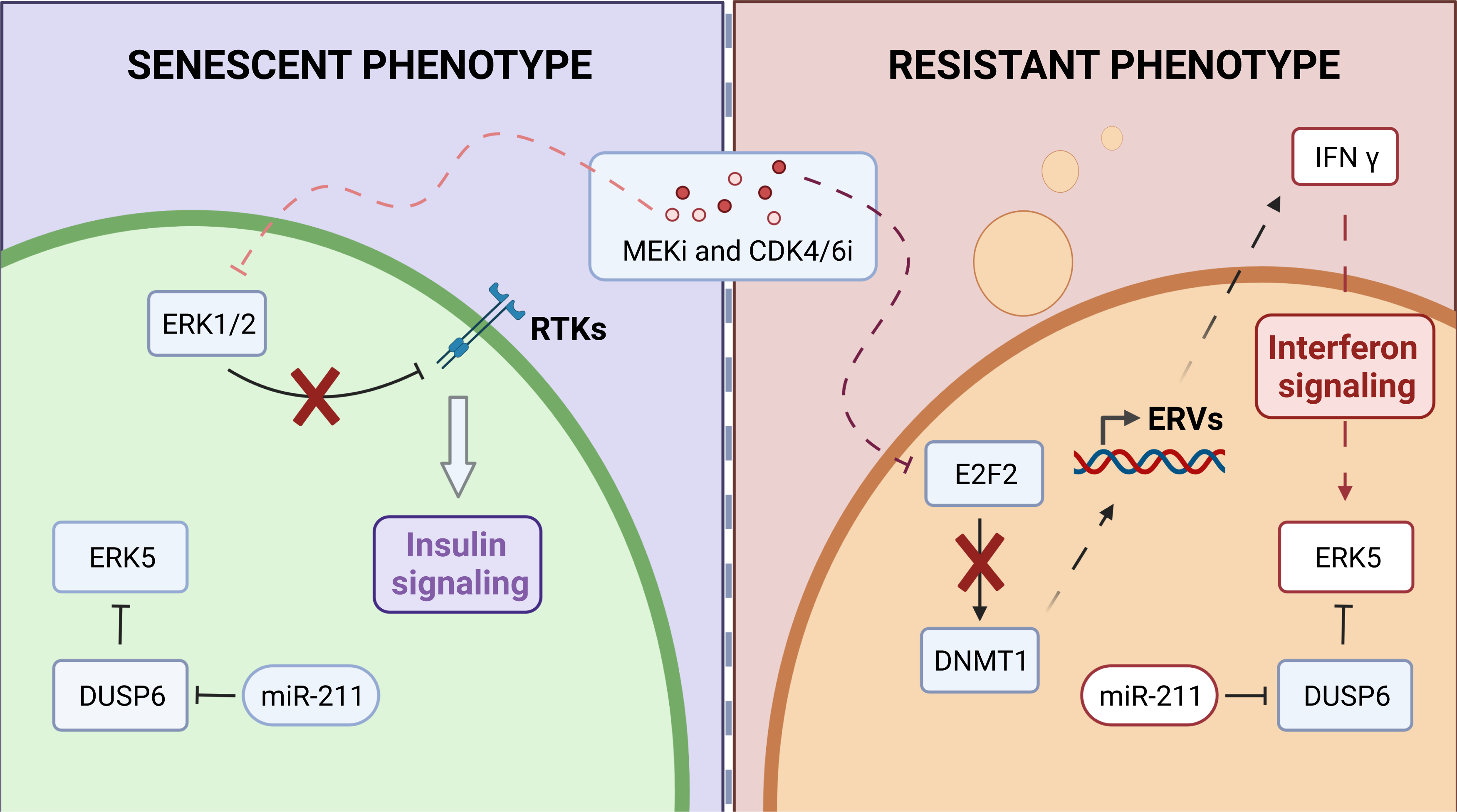
Graphical summary. The phenotypes observed across three different NRAS mutant melanoma cell lines can be summarised as “senescent” (left) and “resistant” (right). Insulin signalling is associated with the “senescent” phenotype and can be explained by the mechanism of action of MEKi, which downregulates ERK1/2, suppressing negative feedback on RTKs. Interferon signaling Can be explained by CDK4/6i, which downregulates DNMT1 leadingand lead to hypomethylation of the genome and expression of Endogenous Retroviruses (ERVs). Induced expression of intracellular ERVs trigger an innate immune response and production of interferons. IFN gamaγ can activate ERK5, inducing EMT. This phenotype is moreover associated with upregulation of miR-211-5p, which indirectly regulates ERK5.

## Methods

### Cell culture

NRAS-mutant MELJUSO, SKMEL30 and IPC298 (ACC-74, DSMZ; ACC-151, DSMZ; ACC-74, DSMZ) and BRAF-mutant A375 and 624MEL (CRCL-1619, ATCC; kindly provided by Dr. Ruth Halaban, Yale School of Medicine) melanoma cell lines were grown in RPMI-1640 medium supplemented with 10% heat-inactivated fetal bovine serum, L-glutamine (2 mM), streptomycin (100 µg/mL) and penicillin (100 units/mL). The cells were kept at 37°C, in a humidified atmosphere with 5% CO2 and sub-cultivated twice a week. Resistance of MELJUSO, SKMEL30, and IPC298 cells to the MEK and CDK4 inhibitors was induced by treatment with 110nM, 35nM or 16nM of Binimetinib, respectively, and 1μM of Palbociclib for 33 days.

### Total RNAseq

Transcriptome sequencing was performed using total RNA. Libraries were prepared using “TruSeq Stranded Total RNA Library Prep Gold” Kit (Illumina, USA) and sequenced as 2x75bp on a NextSeq 550 (Illumina, USA). The sequencing was carried out at Luxgen platform, LNS. Data were aligned to the reference genome (hg38) using STAR (v 2.7.9a) and differential expression analysis was performed using R (v 4.2.2) and DESeq2 (v 1.34.0). This analysis was automated in a snakemake workflow (v 0.3.2) [72] (https://gitlab.lcsb.uni.lu/aurelien.ginolhac/snakemake-rna-seq) using docker file (v 0.6) (https://hub.docker.com/r/ginolhac/snake-rna-seq/tags). Mixing all cell lines into one analysis masked the differences of each time course. Therefore, a subset per cell line was created and the differential gene expression was performed independently. To predict senescence scores with SENESCopedia [18], data were reprocessed and aligned to GENCODE (v 34) using kallisto (v 0.46.2) according to the recommendations of the authors.

### Gene-list based pathway enrichment

Over-Representation Analysis (ORA) and Gene Set Enrichment Analysis (GSEA) were performed using R (v 4.2.2) and the package “clusterProfiler” (v 4.0.5). The Hallmark Gene Set Collection from MSigDB were used as reference. Differentially expressed genes used for GSEA were ranked according to the π value, which takes into consideration both the statistical significance and the biological relevance through the product of the p-value and the fold change [73]. The results were represented using the package “pheatmap” (v 1.0.12) using the default “Ward.D2” clustering algorithm. ORA results were displayed using the “ggplot2” (v 3.3.6) package.

### Kinase activity (PamGene)

To assess the kinase activity within the NRAS mutant melanoma cell lines MELJUSO and SKMEL30 to a combination of CDK4/6 and MEK inhibitors, cell samples were collected at day 0, 1, 4 and 33. Cell pellets were lysed in Mper (78501, Thermo Fisher) supplemented with Halt Phosphatase Inhibitor Cocktail (78428, Thermo Fisher) and Halt Protease Inhibitor Cocktail, EDTA free (78437, Thermo Fisher), for 15 min on ice. The lysate was clarified (10000g, 15 min, 4°C) before aliquoting and freezing at -80°C. Adopting a balanced design, three replicates for each condition and cell line were assessed for both tyrosine kinase activity on PTK chips (86402, PamGene International B.V., ’s-Hertogenbosch, the Netherlands) and serine threonine kinase activity on STK chips (87102, PamGene International B.V., ’s-Hertogenbosch, the Netherlands) using the manufacturer’s protocol. Each chip contains a panel of 196 (PTK) and 144 (STK) phorylatable peptides. The phosphorylation of these peptides by native kinases in the applied lysate was quantified by sequence agnostic fluorescently coupled anti phosphorylation antibodies (Suppl. Fig.4). Image processing was achieved using the instrument manufacturer’s Bionavigator software before applying COMBAT normalization and performing Upstream Kinases Analysis (UKA) using a multiple hypothesis testing approach. This identified responsible kinases for the observed phosphorylation status with a median final score for the likelihood of the kinase being responsible for observed phosphorylation to which a cutoff of 1.2 was applied and a median kinase statistic indicating the degree of and direction of kinase activity change between compared conditions. The results were compiled to construct representative kinome trees with an adaptation of the CORAL tool [74] in R (v 4.2.0).

### Network-based pathway enrichment

To identify biological pathways enriched in a protein list, we applied a network-based protein set enrichment analysis approach using EnrichNet [75]. The inputs are a list of the PamGene protein identifiers and the database of interest from which reference protein sets will be extracted. The databases include (KEGG, BioCarta, WikiPathways, Reactome, PID, and GO. Using a genome-scale molecular interaction network, we map the target and reference datasets and then perform a network analysis, which involves two steps: i) the use of a random walk with restart (RWR) algorithm to score the distance between the mapped target protein set and reference datasets in the network. ii) the comparison of the calculated scores against a background model. Extensive information on how the RWR method for node relevance scoring was developed is found in [75]. A network similarity score (XD-score) measures how close the protein set of the PamGene data is to the pathways in the databases. The XD score is compared to a classical enrichment analysis score (p-values) which measure the significance of the overlap between pathways and the protein set. A similarity ranking table of the reference datasets is generated, including their network-based association scores (measured using the XD-score). The table ranks the pathways from a chosen database to the protein set of the PamGene data. Using an interactive graph-based visualization, we generate interoperable interaction networks for each pathway. This network (XML-based) highlights the proteins that overlap in pathways, illustrating their interactions.

### Small RNAseq

Small RNAseq was performed on the same samples used for total RNAseq. Libraries were prepared using “QIASeq miRNA Library kit” (Qiagen, Germany) and sequenced as 2x75bp on a NextSeq 550 (Illumina, USA). The sequencing was carried out at Luxgen platform, LNS. Data were aligned to the miRBase (v 2.2.2.1) and piRBase (regulatoryrna.org, v 2.0) using bowtie2 (v 2.3.4.1) and differential expression analysis was performed using R (v 4.2.2) and DESeq2 (v 1.34.0). This analysis was performed using a snakemake workflow adapted from a previous workflow designed by https://github.com/jounikuj/ for this library preparation. Briefly, the snakemake workflow applies 4 passes of trimming (including UMIs) and map the obtained reads on miRBase using bowtie2 with the parameter –very-sensitive-local. Next, the corresponding un-mapped reads are aligned to piRBase and the successfully mapped read coming from these 2 passes of alignment are merged in a unique BAM file per sample. Finally reads are counted, after UMI collapsing (UMI_tools v. 1.0.0) using a combination of samtools (v 1.9) and bash commands.

### qCLASH

qCLASH sample preparations were performed exactly as previously described by Kozar *et al.* [16] Briefly, 50 million cells were UV cross-linked at a wavelength of 250nm and 600J/cm2. AGO protein immunoprecipitation was performed using Protein G Dynabeads (Invitrogen), and 10μl 2A8 anti-Ago antibody (Millipore) per sample. qCLASH libraries were sequenced on a NextSeq500 instrument with a read length of 85bp. The raw sequences were pre-processed and analyzed as previously described [16] using the bioinformatics pipeline *hyb* [76], and custom scripts available at GitHub (https://github.com/RenneLab/qCLASH-Analysis).

### miRNA network construction and representation

qCLASH data were analysed using R (v 4.2.2) and the “tidyverse” package. Interaction lists were converted to “igraph” objects with the corresponding package (v 1.3.4) and further exported to cytoscape using “RCy3” (v 2.12.4). Final representations were made using Cytoscape (v 3.8.2) and the plugin “Legend Creator” (v 1.1.6).

### RT-qPCR

RNA was extracted from cell lysates using the Quick-RNA MiniPrep kit with on-column DNase I treatment (Zymo Research, USA) according to the mnufacturer’s protocol. RNA purity and quality were assessed using the NanoDrop2000 Spectrophotometer (Thermo Scientific, USA). 500ng of RNA were used to perform reverse transcription using the miScript RT II Kit and HiFlex Buffer Reverse Transcription (Qiagen, Netherlands). Quantitative polymerase chain reaction (qPCR) was performed using the CFX384 Detection System (BioRad, USA) using 5ng and 50ng of cDNA as a template for subsequent miRNA and mRNA expression detection, respectively. Primers were purchased from Qiagen (miScript Primer Assay range) and from Eurogentec (primer list Supp. Table 4) to respectively amplify miRNA and mRNA. Amplification curves were analyzed using CFX software (BioRad, USA) and expression was normalised using three housekeeping genes (RNU1A, RNU5A, SCARNA17 for miRNA and HPRT, PPIA, TBP for mRNA).

### Single cell RNA-seq analysis

Single cell RNA-seq data from [Randic et al, in revision] were reprocessed using Seurat (v 4.2.0) and SeuratObject (v 4.1.2). Reads were aligned to human genome hg38 using STAR aligner. Transcripts were filtered to be detected in at least 3 cells and to totalise at least 100 counts across cells. Cells were filtered based on mitochondrial transcripts (<=25%), ribosomal transcripts (<= 1%), and a cell complexity (log10GenesPerUMI >=80%). Thresholds for Unique Molecular Identifiers (UMI), genes detected were different between cells lines (for MELJUSO and SKMEL30: 150 and 100, for IPC298: 350 and 150). Finally, mitochondrial, ribosomal and hemoglobin genes were removed, and genes were filtered to have at least 10 counts across the filtered cells.

### iCell data-fusion framework

iCell methodology [15] was applied separately to each of the three cell lines (MELJUSO, SKMEL30 and IPC298) at day 0, day 4 and day 33, to obtain integrated networks that encompass the condition-specific (i.e., defined by a cell line and day) PPI, COEX, and single-cell RNA-seq data from our lab [Randic et al, unpublished]. We used the iCell methodology because it is an intermediate data integration method that uses Non-negative Matrix Tri-factorization (NMTF) to simultaneously decompose multiple relational matrices through the inference of a single joint model [15, 77].

Therefore, it does not suffer from information loss of early integration approaches that combine all datasets into a single dataset before integration, or late integration methods that first build models for each dataset and then combine them into an integrated model. Additionally, NMTF-based methods are co-clustering methods that encode the high-dimensional input data into three low-dimensional matrix factors [78]. The clustering information is contained in the so-called cluster indicator matrices that are easily interpretable (unlike the resulting matrices of other artificial intelligence algorithms), making them ideal for mining biological data.

For each condition (treated cell lines and time points), we constructed condition-specific Protein-Protein Interaction (PPI) and gene co-expression (COEXP) networks as follows. We collected the experimentally validated protein-protein interactions of *Homo sapiens* from BioGRID (v 4.3.195) [32], from which we excluded genes (nodes in the network) that are not expressed in our condition-specific bulk RNA-seq data (zFPKM < −3, [79]). To generate the condition-specific COEXP networks, we applied Spearman’s correlation between the gene expression profiles from the condition-specific bulk RNA-seq data, using a 0.9 correlation threshold with a *p-value ≤ 0.01*. These two condition-specific networks were further filtered to exclude genes (nodes in the networks) that are not expressed in any single cell according to the corresponding condition-specific single cell RNA-seq data.

For each condition, we followed the iCell methodology [15] to integrate the condition-specific PPI, COEXP, and single-cell expression data together. To this aim, we used Non-negative Matrix Tri-Factorization to simultaneously decompose the adjacency matrices of the PPI and COEXP networks, *A_1_* and *A_2_*, respectively, and the single cell RNA-seq expression data, *E*. More precisely, the two adjacency matrices, *A_1_*and *A_2_*, are simultaneously decomposed as the products of three matrix factors each, *A_1_* as the product of matrix factors *G_1_*, *S_1_* and 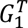, and *A_2_* as the product of matrix factors *G_1_*, *S_2_* and 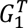, i.e.: 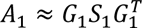 and 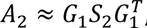, where *G_1_* is interpreted as the cluster indicator matrix of the genes and matrices *S_1_*and *S_2_* are interpreted as the compressed representations of the PPI and COEXP networks, respectively. The condition-specific single-cell RNA-seq expression matrix, *E*, is decomposed into the product of three matrix factors *G_1_*, *S_3_* and 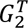 as: 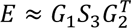, where *S_3_* is interpreted as the compressed representation of the single-cell expression data, *G_2_* is the cluster indicator matrix of single cells, and *G_1_* is the cluster indicator matrix of the genes that is shared across the decompositions of all input matrices, which facilitates the information flow and allows for learning from all data.

Furthermore, to ease interpretability, we constrain all matrix factors to be non-negative. We also impose orthogonality constraint to the *G_2_* matrix factor to obtain less ambiguous functional organization captured by the clusters [78].

This decomposition is done by minimizing the following objective function, *F*:

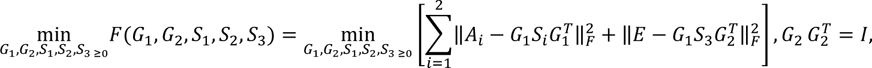

where *‖ ‖_F_* denotes the Frobenius norm and *I* is the identity matrix.

Because minimizing this objective function is an NP-hard continuous optimization problem [80], we heuristically solve it by using a fixed point method that, starting from an initial solution, iteratively applies multiplicative update rules [78] to converge towards a locally optimal solution (described in the next section).

After minimizing *F*, we used the obtained matrix factors to create an integrated network that encompasses the condition-specific PPI, COEXP, and single cell RNA-seq data, which we call iCell network. Following the iCell methodology [15], we created this network by computing 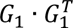 and thresholding the resulting matrix to preserve only the top 1% of the strongest relationships in each row and column.

We apply this methodology for each condition separately, resulting in nine condition-specific iCell networks.

### Fixed point solver

Following the iCell methodology [15], first we derive the Karush-Kuhn-Tucker (KKT) conditions, which are necessary for a solution to be optimal [80]:

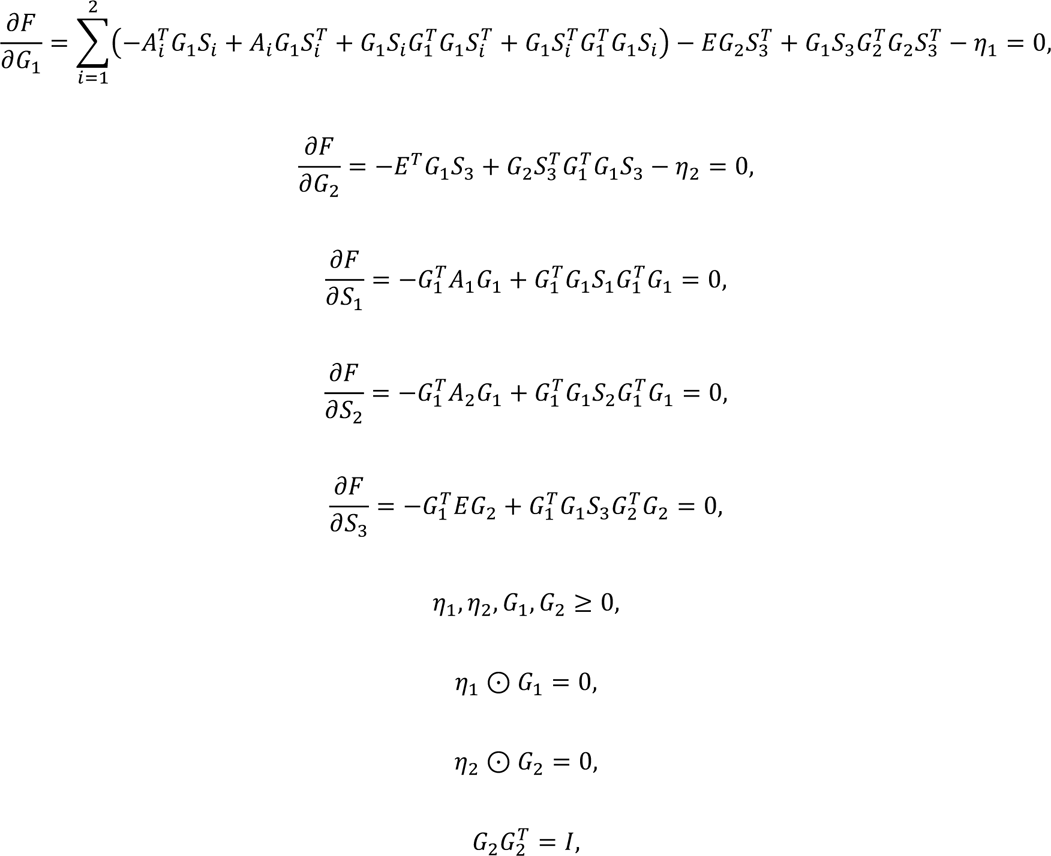

where is the Hadamard (elementwise) product, *I* is the identity matrix, and matrices, *7_1_* and *7_2_*, are the dual variables for the primal constraints *G*_1_ ≥ 0 and *G*_2_ ≥ 0, respectively.

Then, we derive the following multiplicative update rules for all matrix factors, *G_1_*, *G_2_*, *S_1_*, *S_2_*and *S_3_*, to solve the KKT conditions presented above:

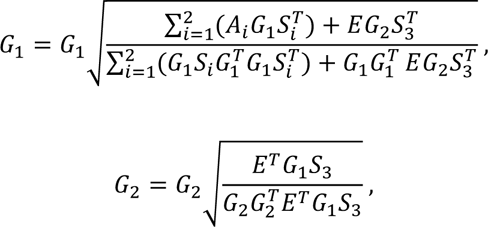

In our fixed point approach, we start from initial solutions of the matrix factors, which we compute using singular value decomposition (SVD) of the original input matrices [81], and iteratively use the derived update rules to compute new matrix factors, *G_1_*, *G_2_*, *S_1_*, *S_2_* and *S_3_*, until convergence. Initializing the matrix factors with SVD reduces the number of iterations that are needed to achieve convergence and makes our solver deterministic [81]. To satisfy the non-negativity constraint of NMTF, we generate the initial solutions by taking the absolute values of the entries from the matrices computed using SVD.

The iterative process ends when the objective function does not decrease anymore, which we measure every ten iterations with: 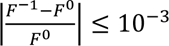, where *F^O^* is the value of the objective function at the current iteration, *i*, and *F^-1^* is the value computed at iteration *i-10*.

### Analyzing the topology of iCells

To mine the biological knowledge hidden in the iCell networks, we applied graphlet-based methods because they are the most sensitive measures of network topology to date [28, 34]. Graphlets are small, connected, nonisomorphic, induced subgraphs of a large network that appear at any frequencies in the network [82]. Within a graphlet, the symmetries between groups of nodes are called automorphism orbits and are used to characterise the different topological positions that a node participates in [82]. These orbits are used to generalised the notion of the node degree (the number of edges that the node touches in the network): for each orbit, the “graphlet degree of the node” in the network is the number of times the node touches (is found at the position of) the particular orbit [28]. Following the methodology of Yaveroğlu *et al*. (2014), we characterised the topology of each node in a network by using the 11 nonredundant orbits of 2-to 4-node graphlets. Therefore, the topology of every node is captured by an 11-dimensional vector called “Graphlet Degree Vector” (GDV).

We compared the global topological dissimilarity between two condition-specific iCell networks by using “Graphlet Correlation Distance” (GCD-11), because it is the most sensitive network distance measure [28]. To do this, first we characterised the global topology of a network with its “Graphlet Correlation Matrix” (GCM), which is an 11 × 11 symmetric matrix that contains the Spearman’s correlations between GDVs over all nodes of the network. Then, the GCD-11 between two networks is the Euclidean distance between the upper triangle values of their GCMs. The lower the GCD-11 value, the closer the topologies of the networks [28].

Additionally, we measure the change in the wiring of a gene between two condition-specific iCell networks by using the “Graphlet Degree Vector Similarity” (GDVS) between the GDVs of the node in the two networks, which is computed as follows. Given the GDV vector of a gene (node) in one network, *a*, and the GDV vector of the same gene (node) in another network, *b*, the distance between the *i^th^*coordinates of vectors *a* and *b* is defined as:

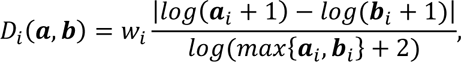

where *w_i_* is the weight of orbit *i* that accounts for dependencies between the orbits [29]. Then, GDVS between vectors *a* and *b* is defined as:

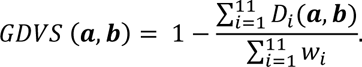

GDVS is the similarity measure in (0,1] range, such that the similarity equal to 1 means that the two GDV vectors are identical, and thus, that the gene has the same local topology in both networks. We use GDVS to identify the genes whose topology changed between two condition-specific iCell networks, which we termed “perturbed” (i.e., whose GDVSs were close to 0), as well as the genes whose local topology did not change, which we termed “stable” (i.e., whose GDVSs were close to 1). These comparisons were made between different time points for the same cell line, as well as between different cell lines for the same time point.

### Clustering

For each cell line and each time point, we performed clustering of the Graphlet Degree Vectors (GDVs). Prior to clustering, the features were weighted by using previously defined weights [29] and then log-transformed. In addition, both the NMTF and GCM matrices were scaled to a 0 mean and unit variance. For each input matrix, we first computed the Hopkins statistics to estimate cluster tendency. Then, to identify the numbers of clusters, we estimate the cluster validity of potential solutions reporting between 50 and 150 clusters, using 21 different metrics. We rank all clustering solutions for each metric. Then, for each input matrix, the number of clusters is selected based on the solution associated with the minimal median rank across all 21 metrics. We then performed hierarchical and k-medoids clustering using a Pearson correlation-based distance and validated the obtained repartitions using silhouette width and the between/within average distance ratio. All scripts are available on the associated Git repository.

### Network representation of gene predictions

Protein-Protein interactions between TP53, a known regulator of senescence, and genes associated with “EMT” and “INTERFERON GAMMA RESPONSE” which have been linked with senescence escape were extracted from BioGRID (v 4.4.217). A network representation was created by using Cytoscape (v 3.9.1). The representation was further refined by using stringApp (v 2.0.0) and clusterMaker2 app (v 2.3.2) and modified by using Inkscape (v 1.2.1).

### Availability of data and materials

Raw data together with processed files were deposited to: PRJEB59603. Code to reproduce the results in this manuscript are available via GitLab repository (https://gitlab.com/Ahmed7emdan/integrative-multi-omics-analysishttps://github.com/).

## Authors’ contributions

SK conceived the study. SK, FT, DP, MT, TR and VG contributed to the experimental design. VG, FT, and DP acquired total RNA-seq and small RNA-seq data and AM, AG and VG processed, analysed and interpreted the data. TR, DP and CM acquired PamGene data, JL processed the data and performed Upstream Kinase Analysis. MO and AH performed network-based pathway enrichment and created xml models to overlay the inferred kinase activities; VG and CA interpreted the results. IK and HH performed qCLASH method, IK processed the data and AG, VG and HH analysed the hybrids and constructed the miRNA network. HH performed qPCR validations. MB processed the scRNA-seq data. NP, NMD, GC and KM conceived the data integration. KM, GC, and VG performed it and analysed the results under the supervision of NMD and NP. LT performed the gene clustering; AB, MB, and VG analyzed the gene clusters. VG, HH, AH, CA, AB and FT contributed to the figures. VG and SK drafted the manuscript with input from all authors. NP, NMD and MO reviewed the manuscript. All authors read and approved the final manuscript.

## Acknowledgements

The authors want to thank L. Sinkonnen for his suggestions regarding the qCLASH method and its extended applications, J.P. Wroblewska for her feedback and her insights regarding the biology of amelanotic melanoma, A. Ginolhac for his help in the analysis of our RNA-seq data and his tips regarding R language. Several *in silico* experiments presented in this paper were carried out using the HPC facilities of the University of Luxembourg, Varrette et al. 2022, https://hpc.uni.lu.

## Supplementary Materials

**Supplementary Figure 1.**
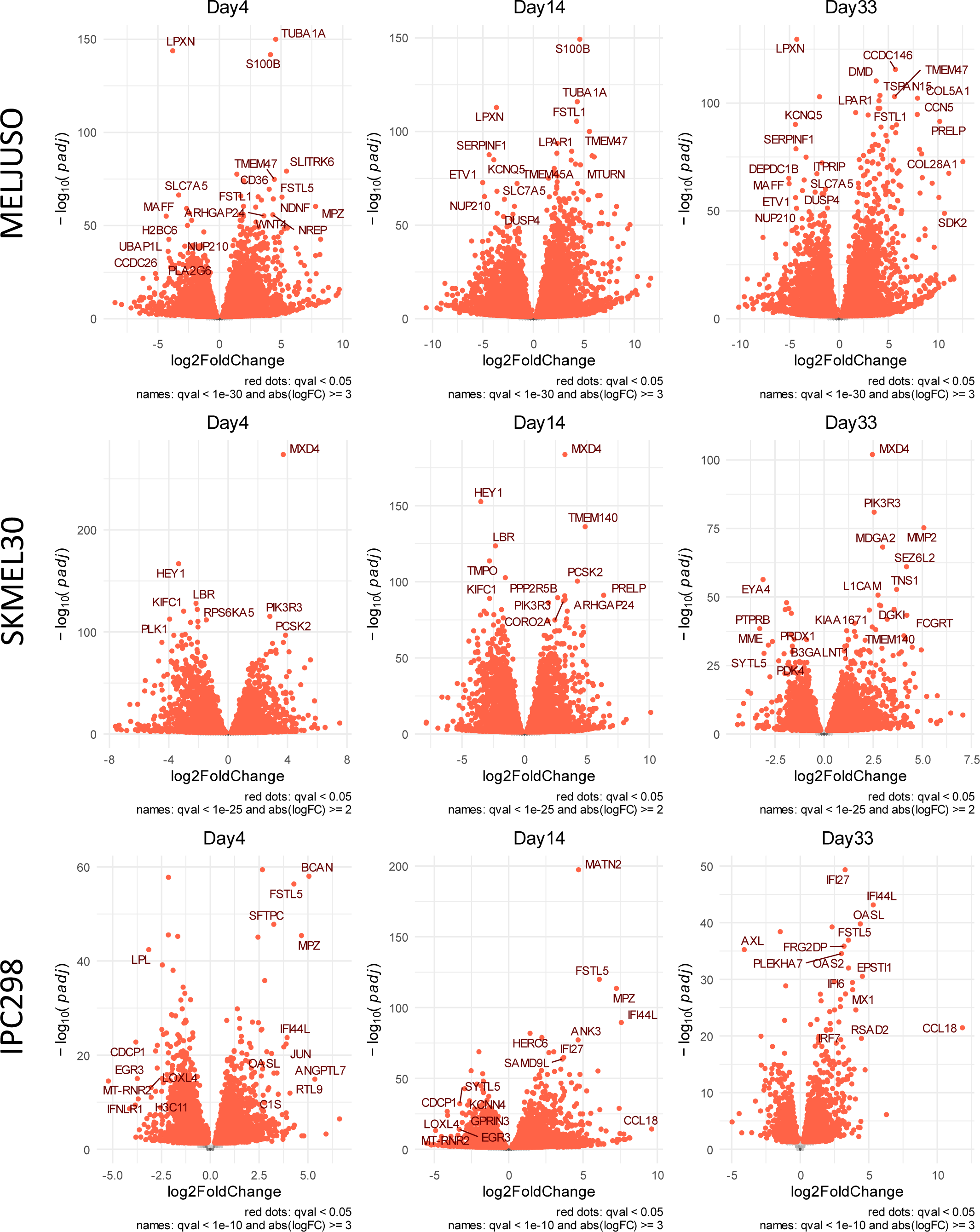
Volcano plots for the total RNA-seq. MELJUSO, SKMEL30 and IPC298 cell lines were treated with CDK4/6i and MEKi.

**Supplementary Figure 2.**
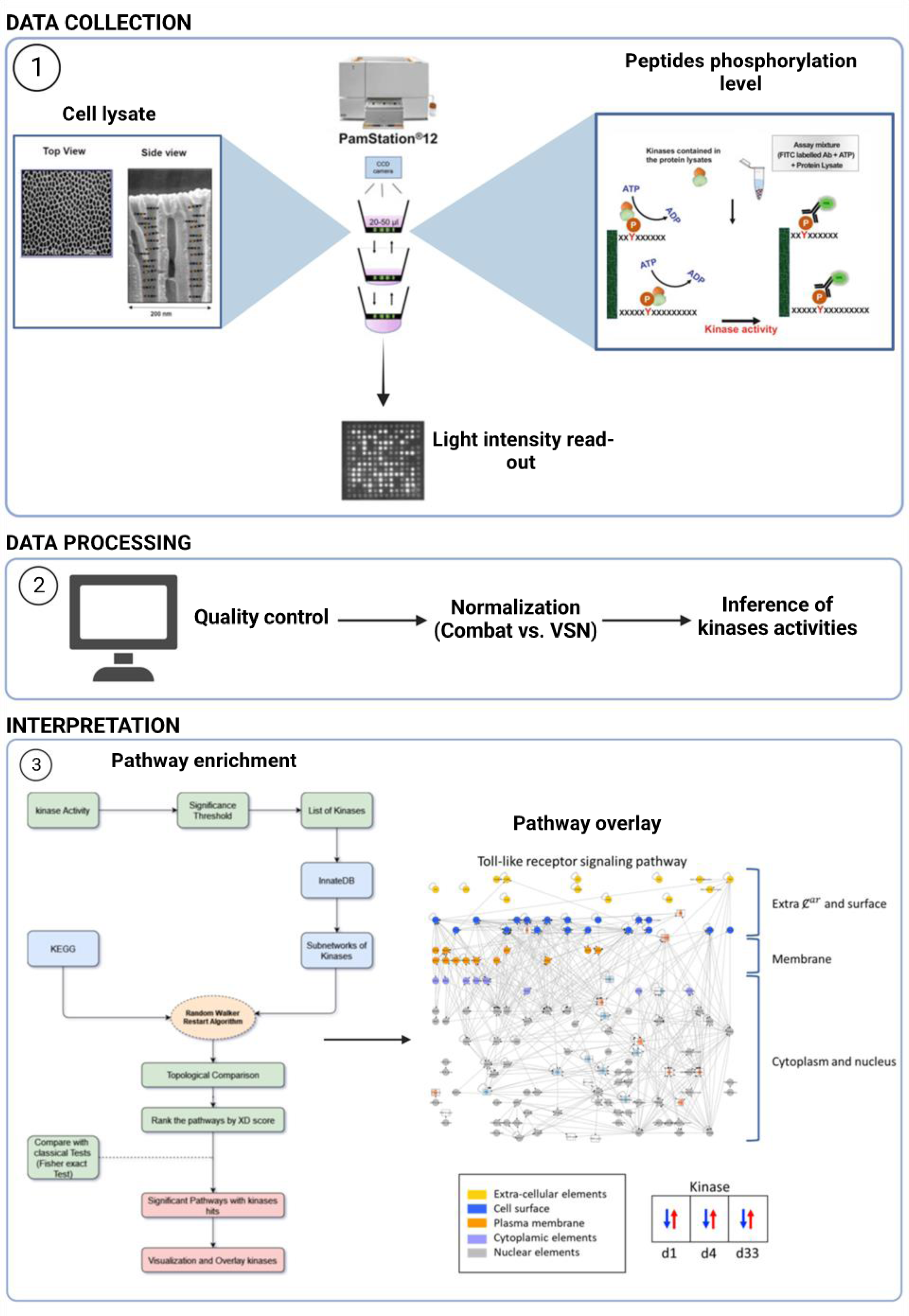
Pamgene data acquisition, processing and interpretation workflow.

**Supplementary Figure 3.**
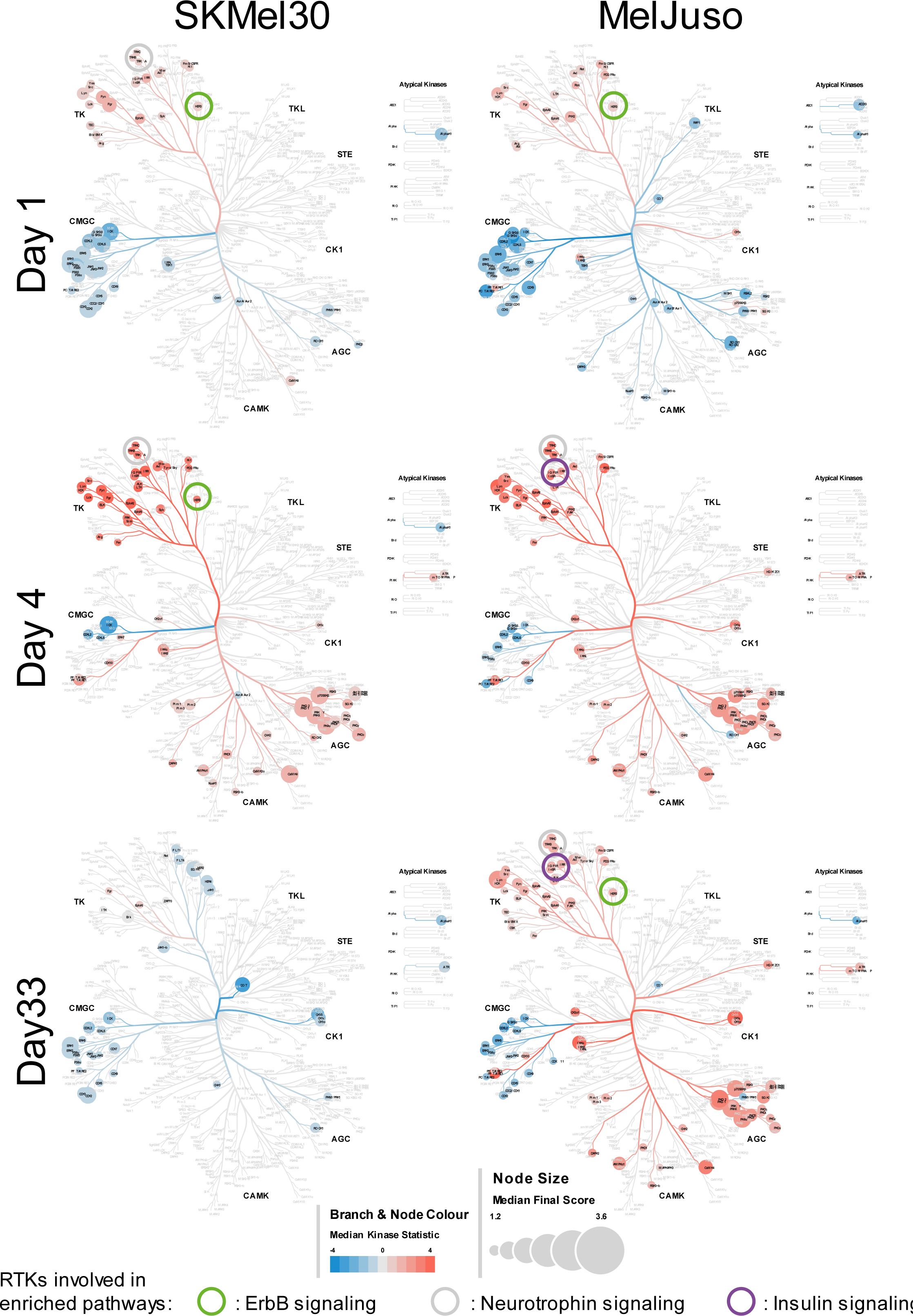
Phylogenetic trees representing the kinome responses to MEKi and CDK4/6i. The size of the leafs represents the “Median Final Score”, a score greater than 1.2 indicates a significant change between conditions. The color of the branchs and leafs shows the “Median Kinase Statistic” which is the difference in kinase activity. Circles highlight RTKs contributing to enriched pathways. SKMEL30 and MELJUSO cell lines show similar responses at early time points but not at 33 days.

**Supplementary Figure 4.**
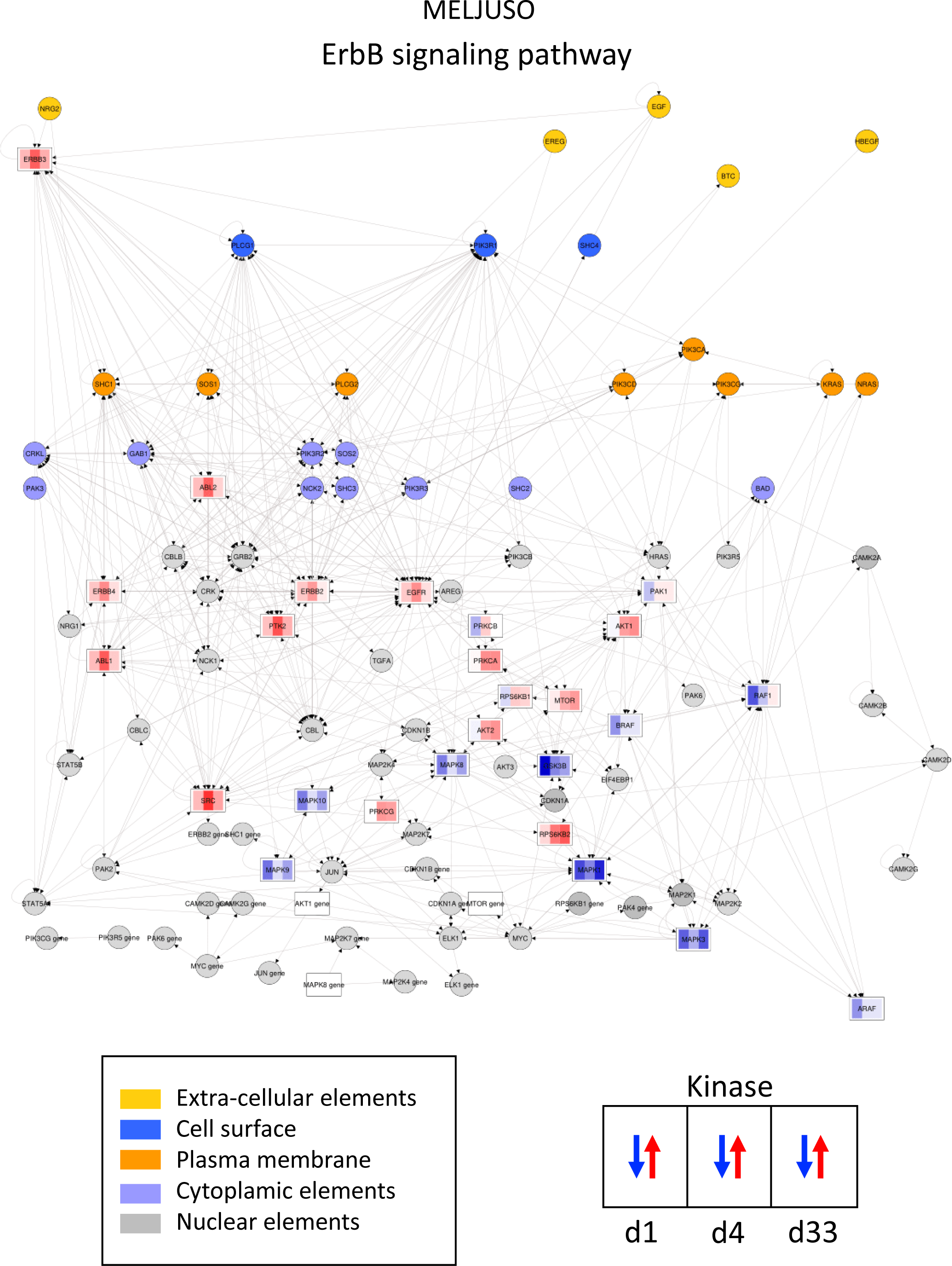
Pathway representation of ErbB signaling in MELJUSO cell line. The color of the nodes represents the cellular localization while kinases are illustrated as rectangles with stripes showing their activity at day 1, day4 and day 33.

**Supplementary Figure 5.**
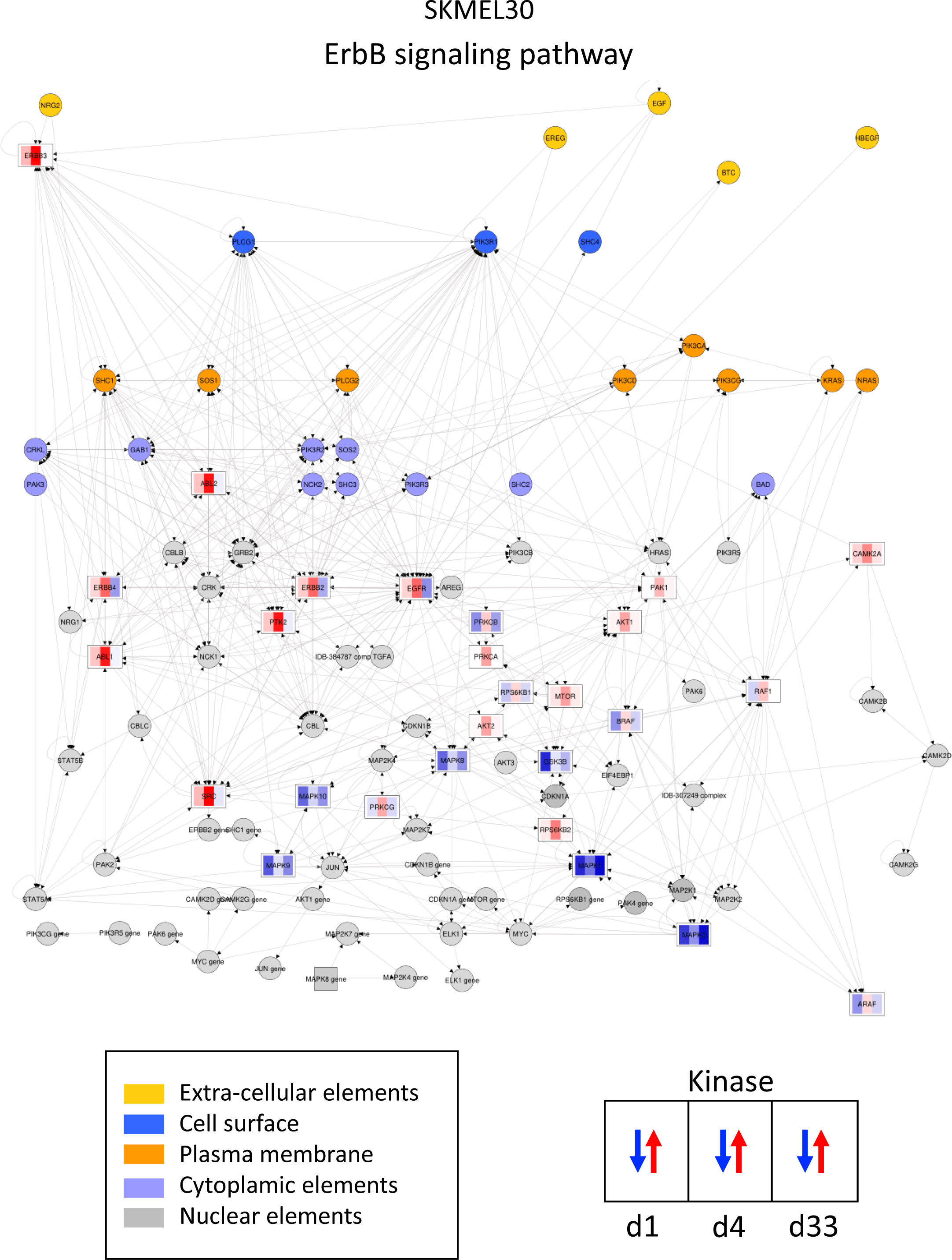
Pathway representation of ErbB signaling in SKMEL30 cell line. The color of the nodes represents the cellular localization while kinases are illustrated as rectangles with stripes showing their activity at day 1, day4 and day 33.

**Supplementary Figure 6.**
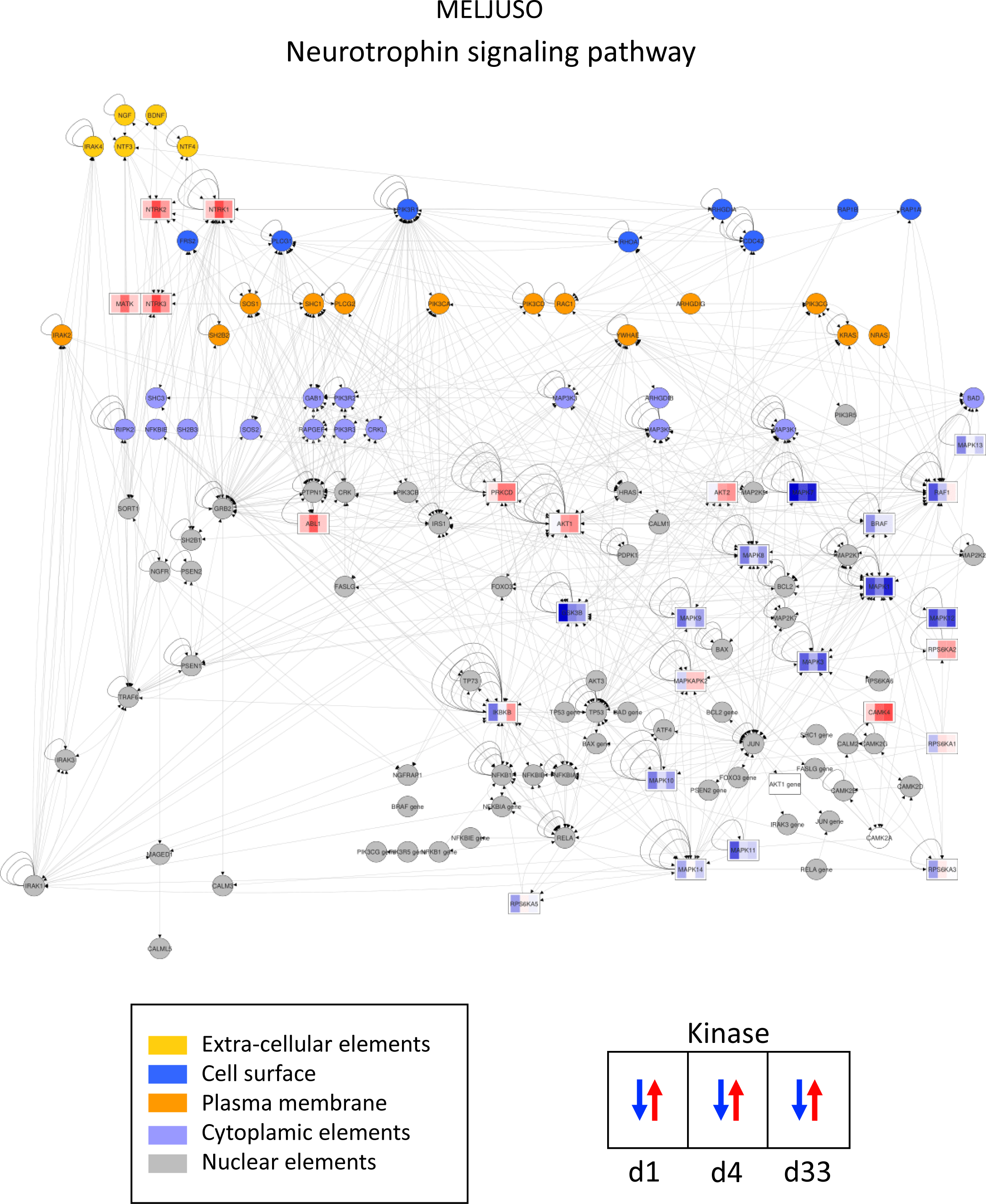
Pathway representation of Neurotrophin signaling in MELJUSO cell line. The color of the nodes represents the cellular localization while kinases are illustrated as rectangles with stripes showing their activity at day 1, day4 and day 33.

**Supplementary Figure 7.**
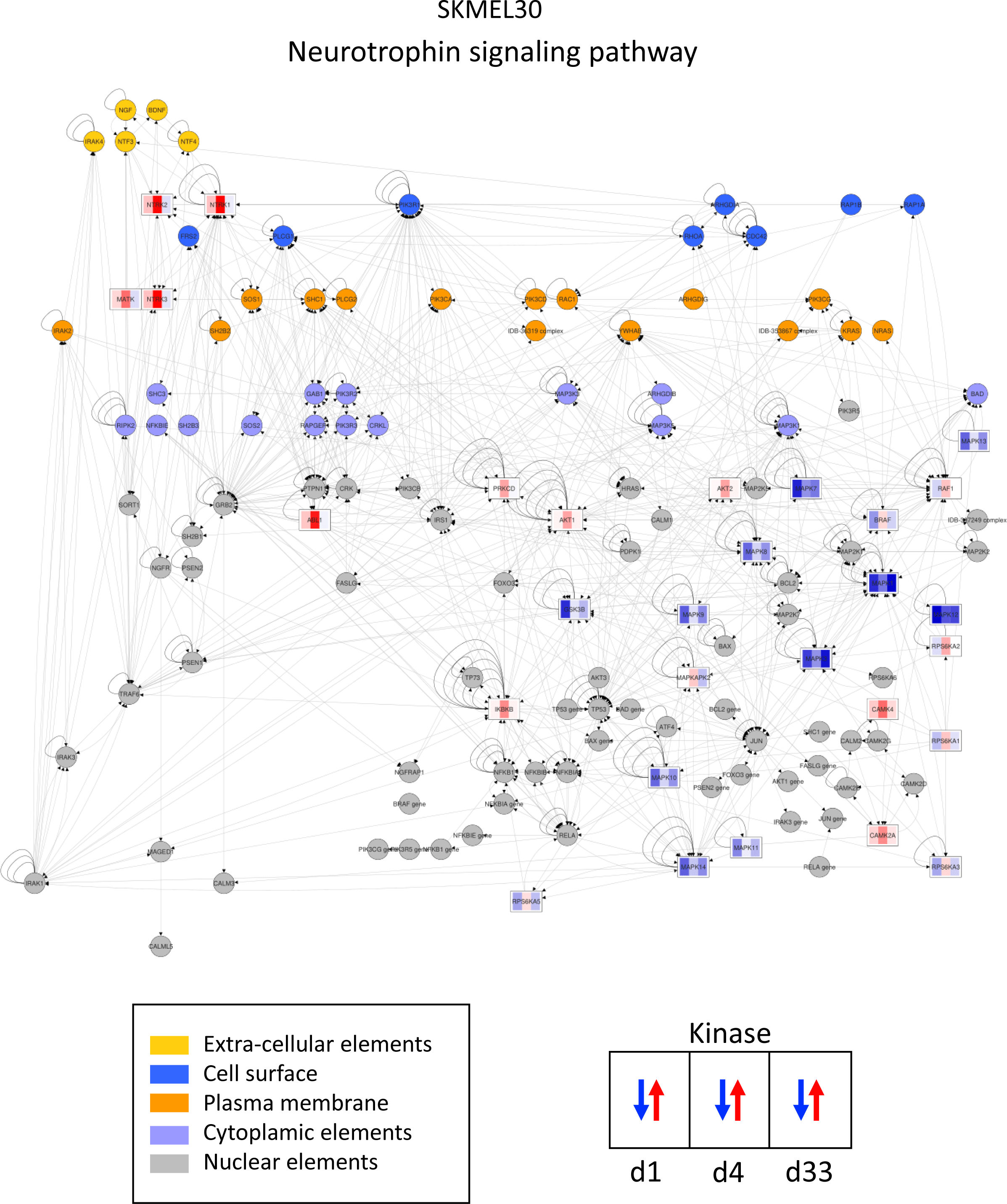
Pathway representation of Neurotrophin signaling in SKMEL30 cell line. The color of the nodes represents the cellular localization while kinases are illustrated as rectangles with stripes showing their activity at day 1, day4 and day 33.

**Supplementary Figure 8.**
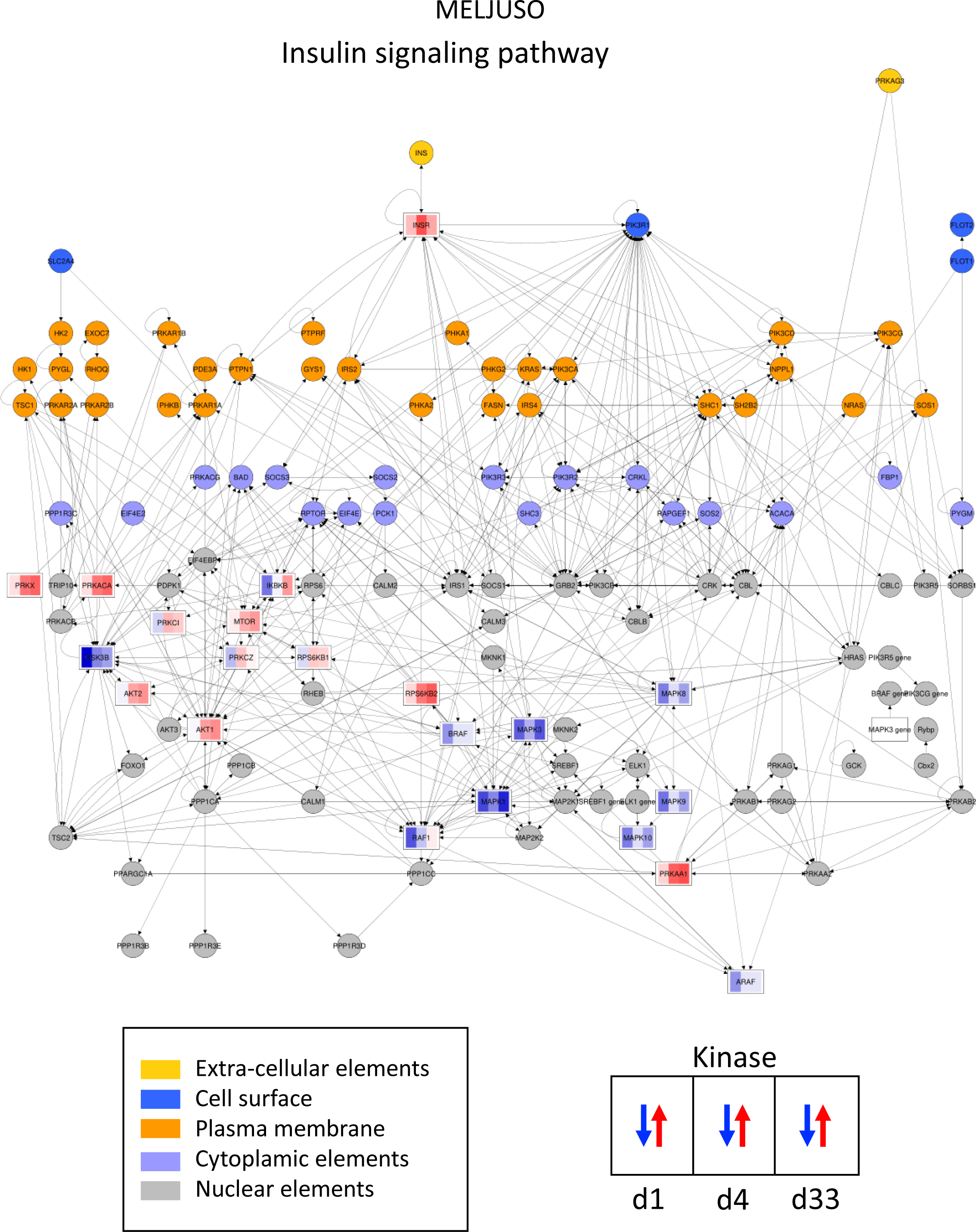
Pathway representation of Insulin signaling in MELJUSO cell line. The color of the nodes represents the cellular localization while kinases are illustrated as rectangles with stripes showing their activity at day 1, day4 and day 33.

**Supplementary Figure 9.**
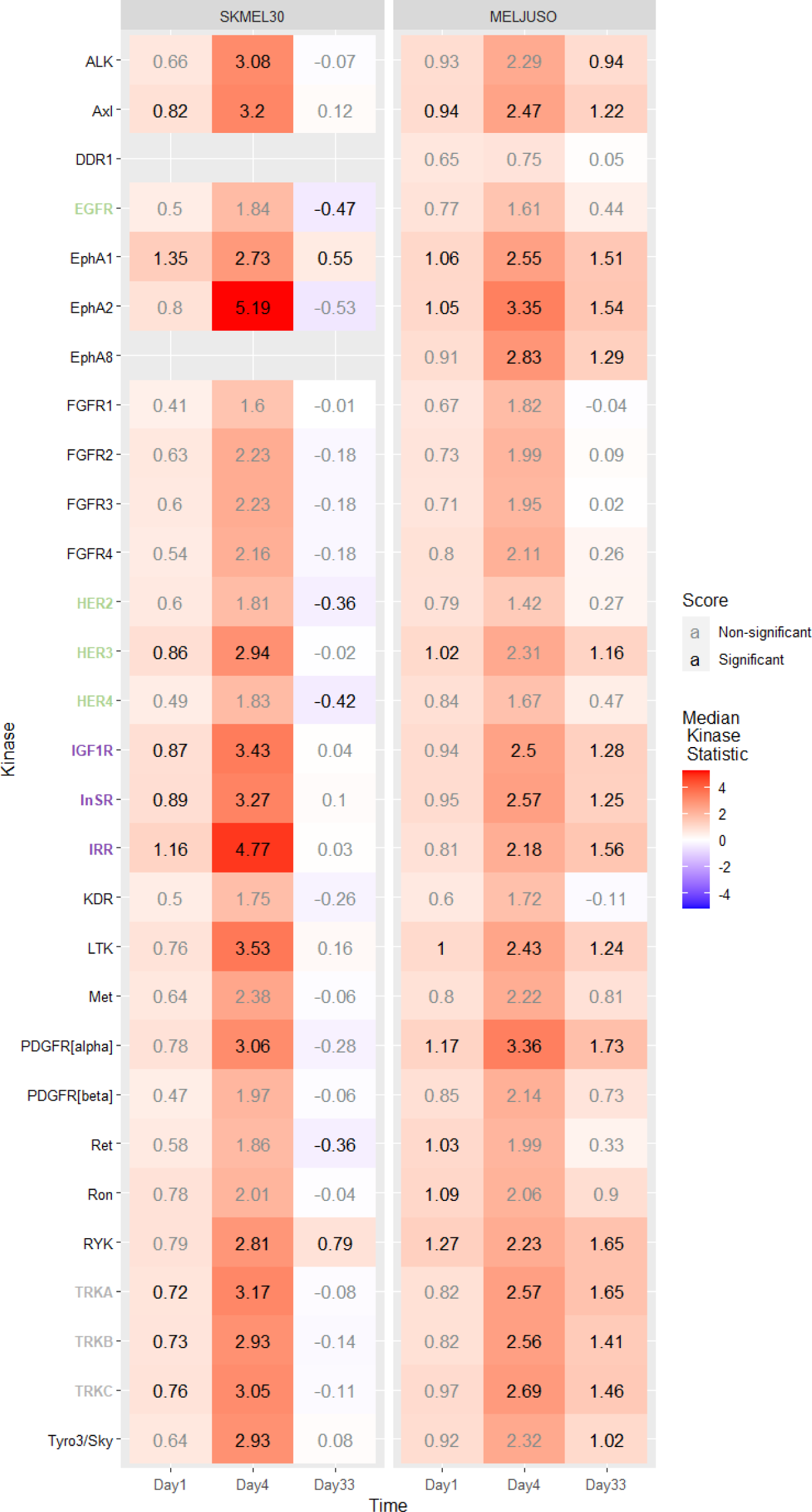
Upregulation of RTKs upon MEKi and CDK4/6i. The heatmap represents the “Median Kinase Statistic” for kinases, significant values are represented in black. RTKs are colored according to the enriched pathways in which they are involved.

**Supplementary Figure 10.**
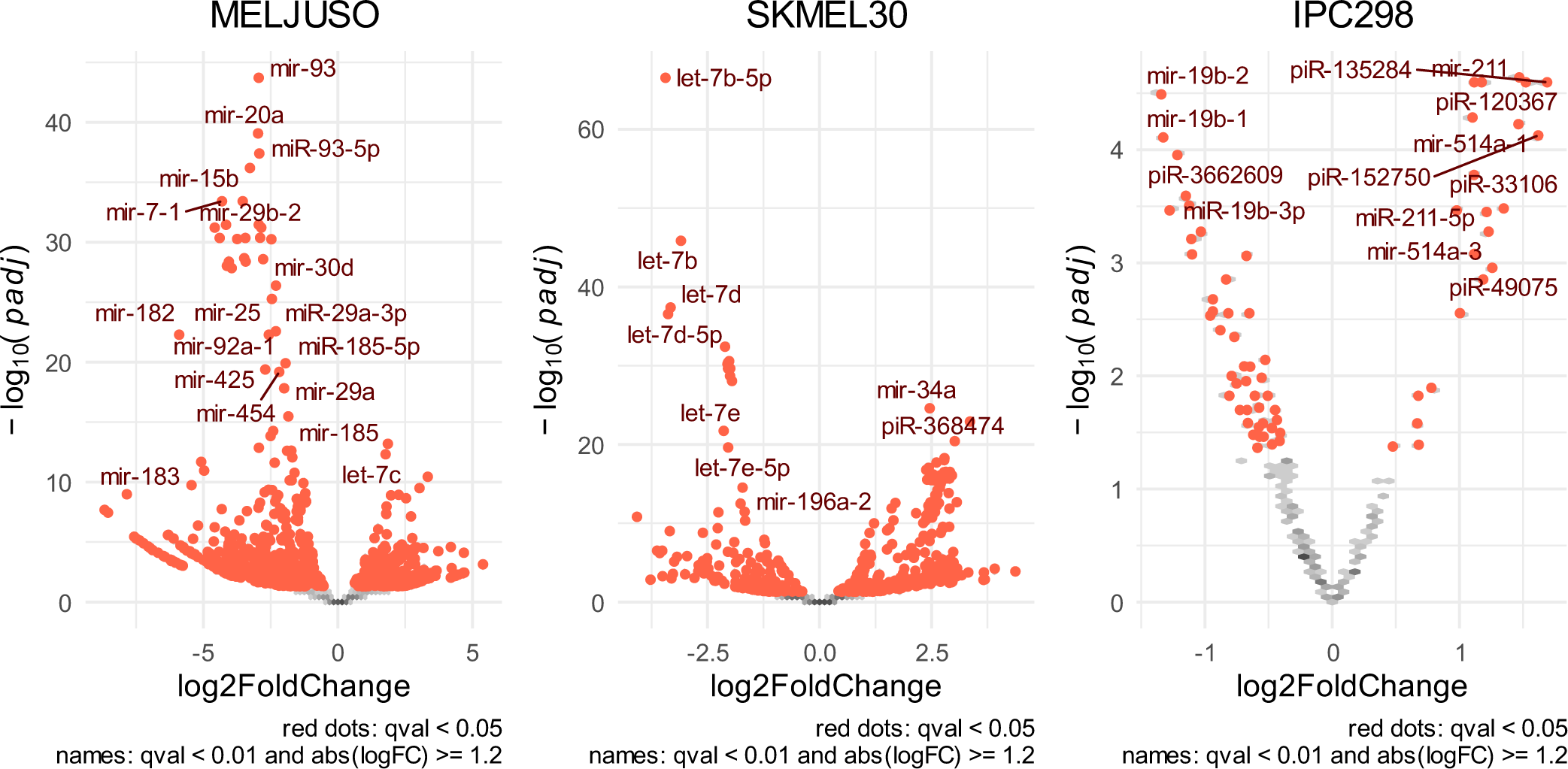
Volcano plots for the small RNA-seq. MELJUSO, SKMEL30 and IPC298 cell lines were treated with CDK4/6i and MEKi.

**Supplementary Figure 11.**
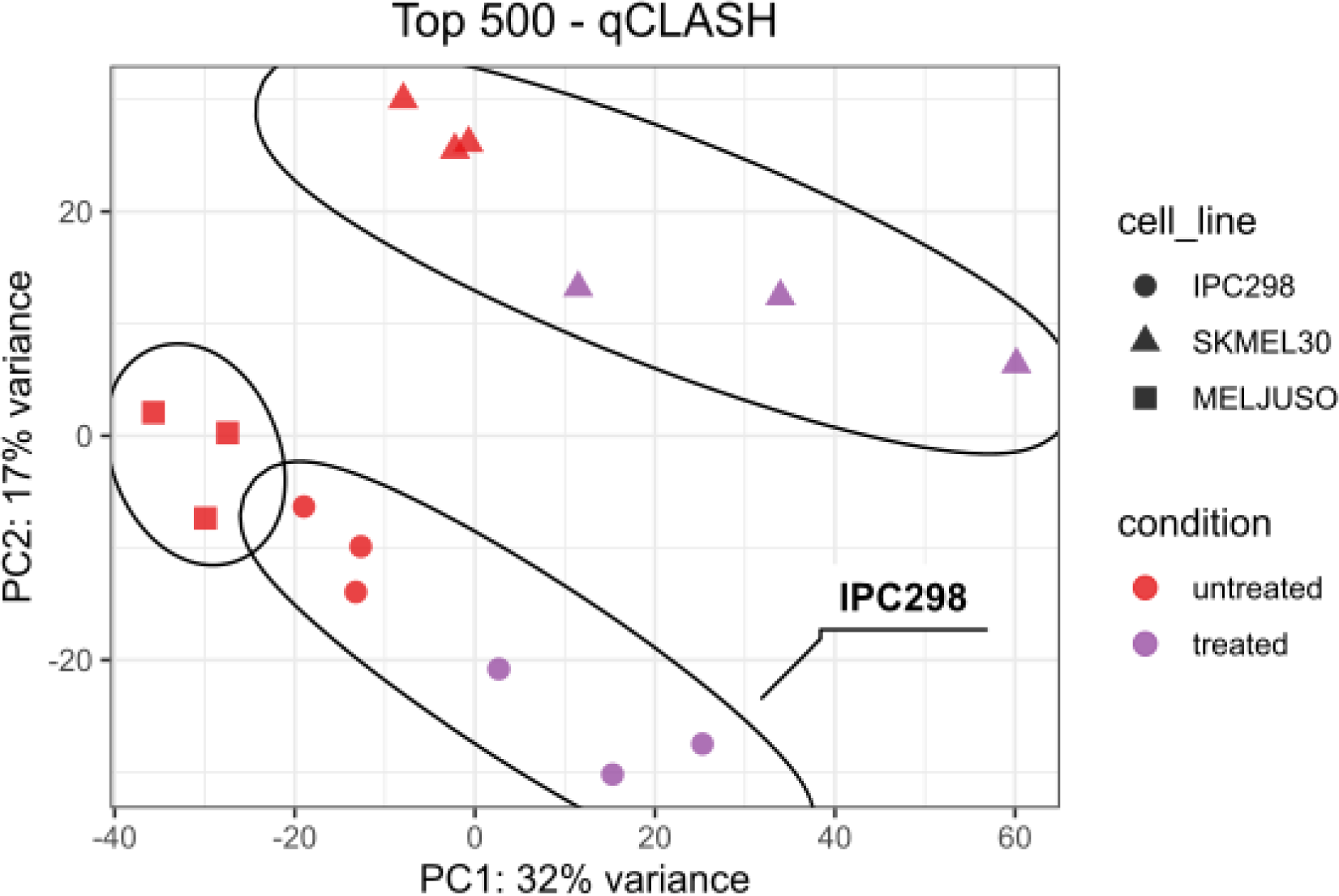
Principal Component analysis (PCA) for the top 500 most detected interactions. PCA clearly distinguishes cell lines and conditions.

**Supplementary Figure 12.**
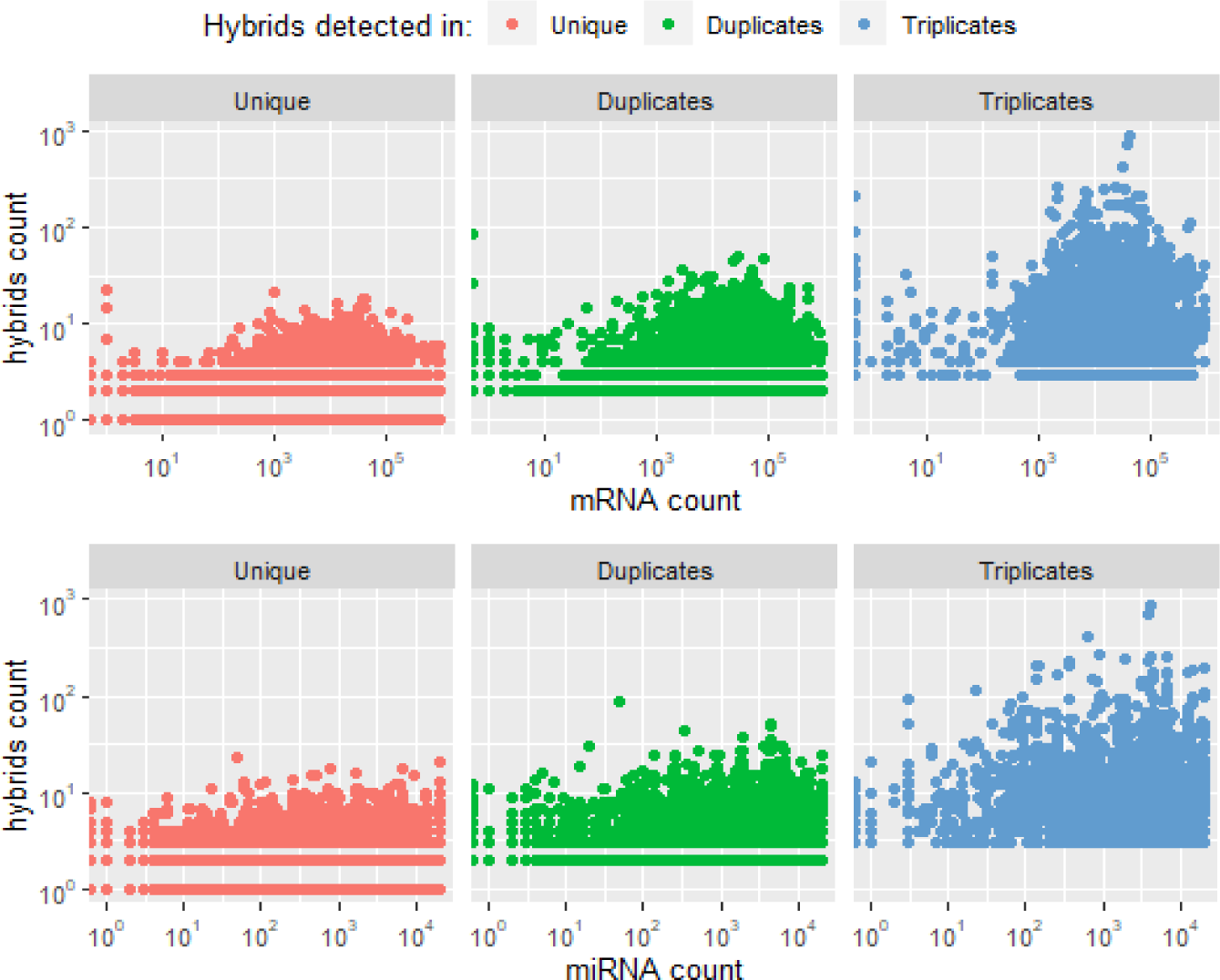
miRNA/mRNA vs hybrids count detected in one, two or three biological replicates. Interaction detected in triplicates are associated with greater expression values.

**Supplementary Figure 13.**
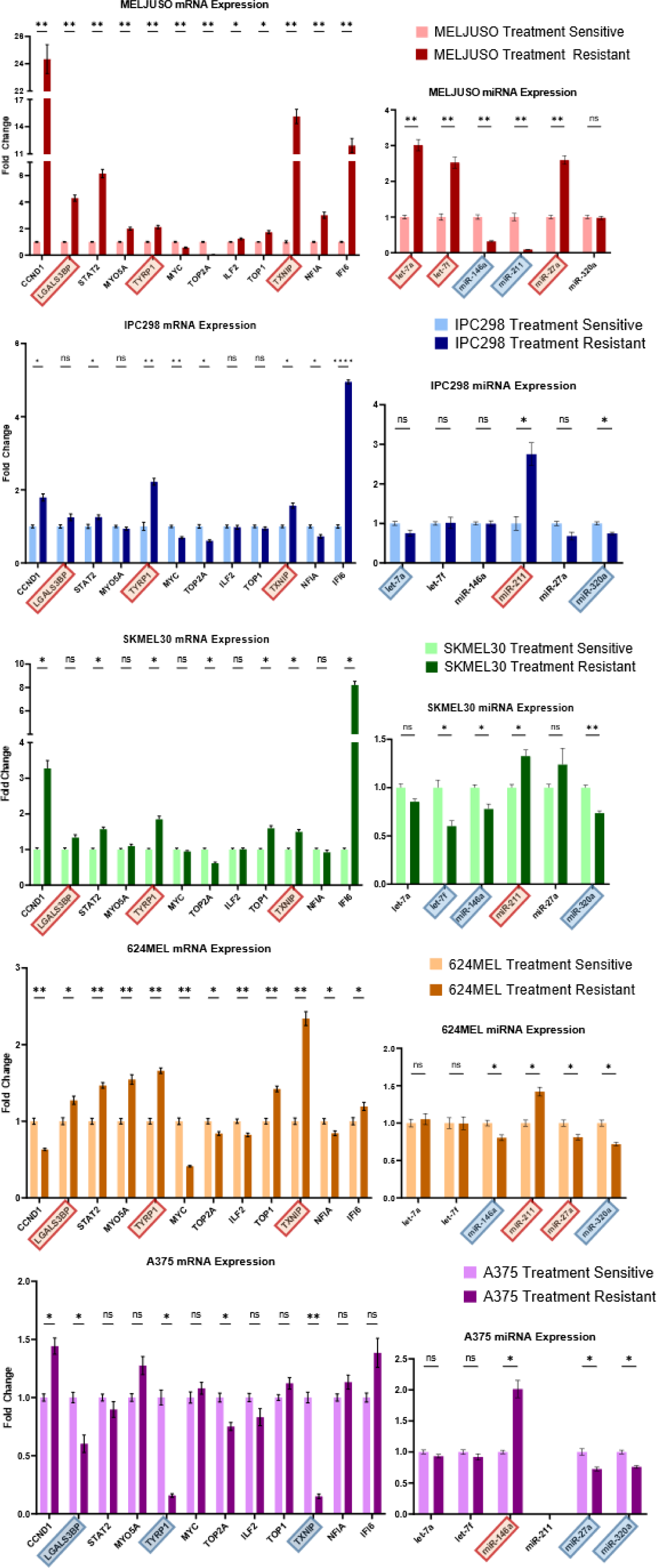
miRNA/mRNA expression detected by qPCR across three NRAS and two BRAF cell lines. Statistical significance was calculated for each condition using multiple welch’s t-test and an FDR of 0.01. Red rectangles highlight upregulated elements and blue rectangles, downregulated elements.

**Supplementary Figure 14.**
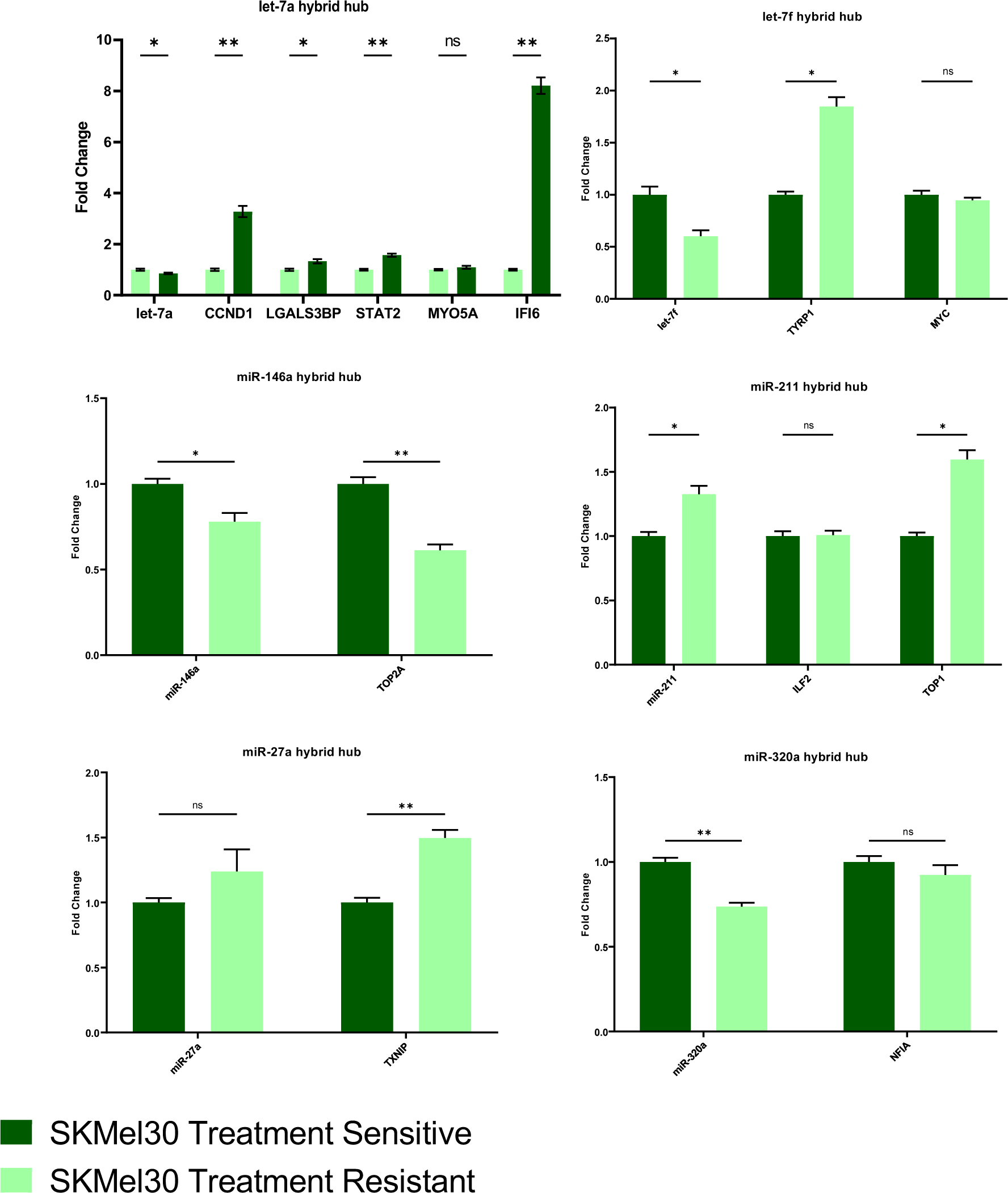
Expression of miRNAs and mRNAs detected by qPCR in let-7a, let-7f, miR-27a, miR-146a, miR-211 and miR-320a hubs in SKMEL30.

**Supplementary Figure 15.**
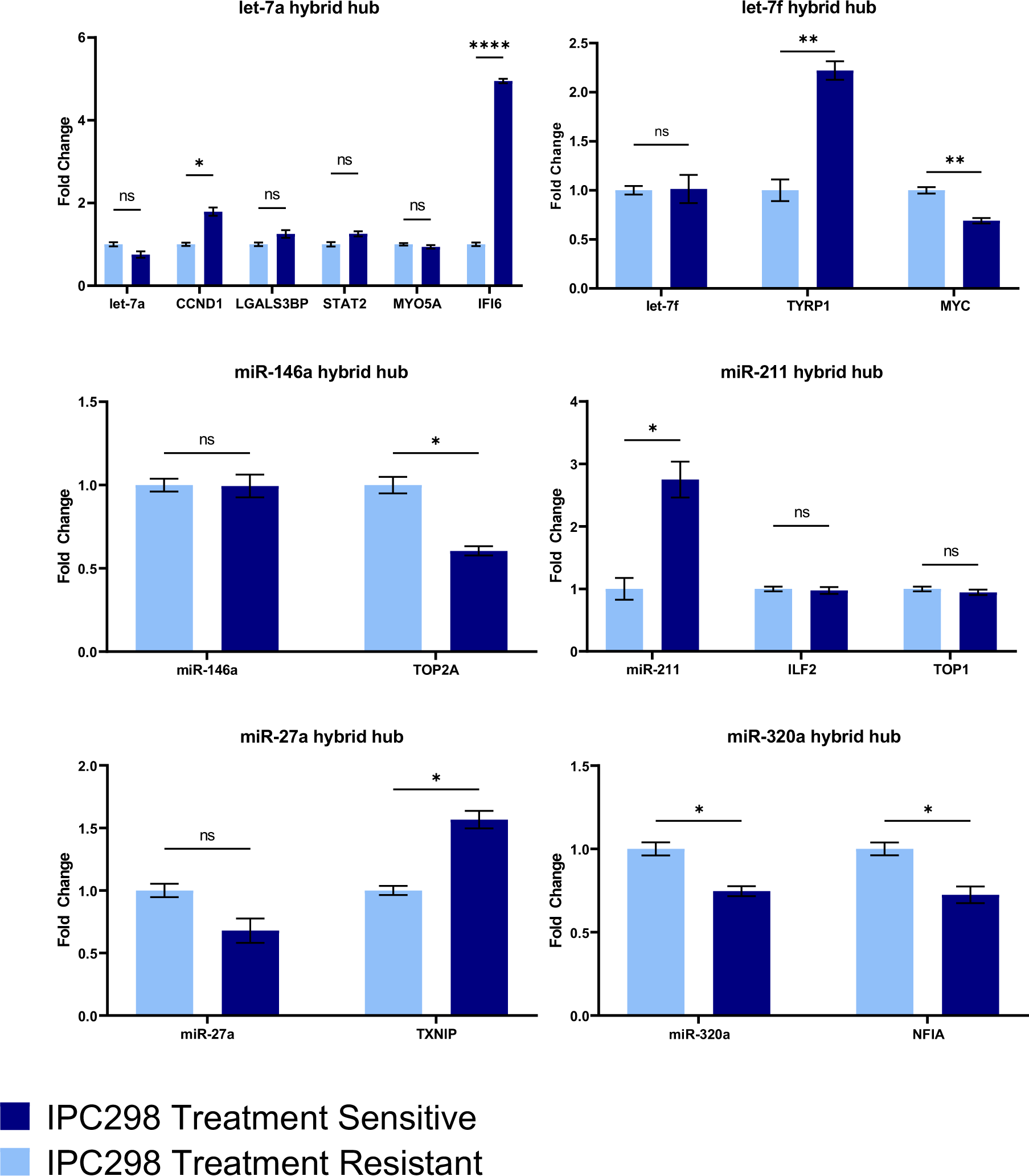
Expression of miRNAs and mRNAs detected by qPCR in let-7a, let-7f, miR-27a, miR-146a, miR-211 and miR-320a hubs in IPC298.

**Supplementary Figure 16.**
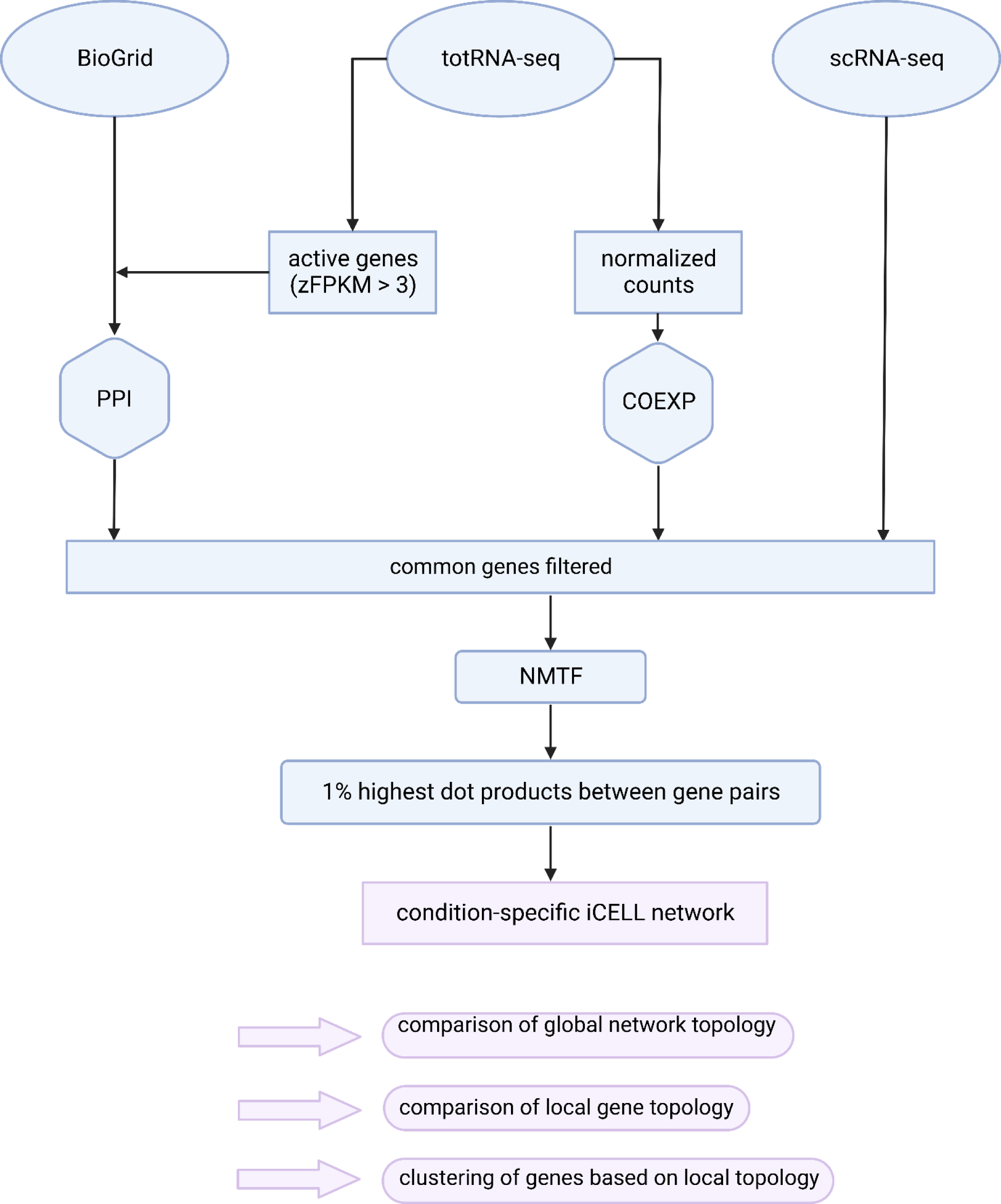
Workflow of the data integration, construction an investigation of the iCELL networks.

**Supplementary Figure 17.**
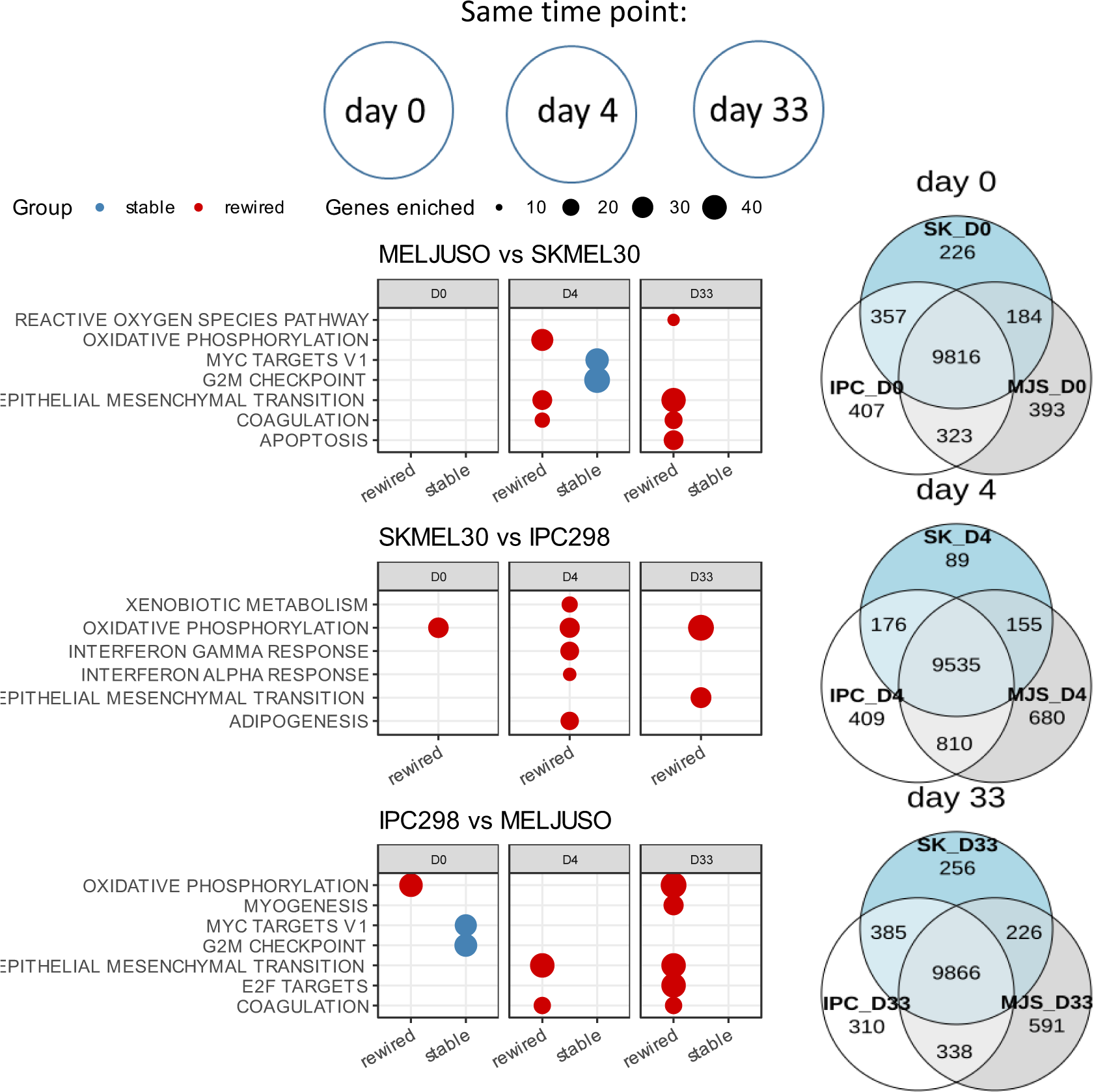
Comparison of gene topology using Graphlet Degree Signature Similarity (GDSS) for matching time points. ORA was performed on the top 10% most “stable” or “perturbated” genes. Venn diagrams represent genes overlap between conditions.

**Supplementary Figure 18.**
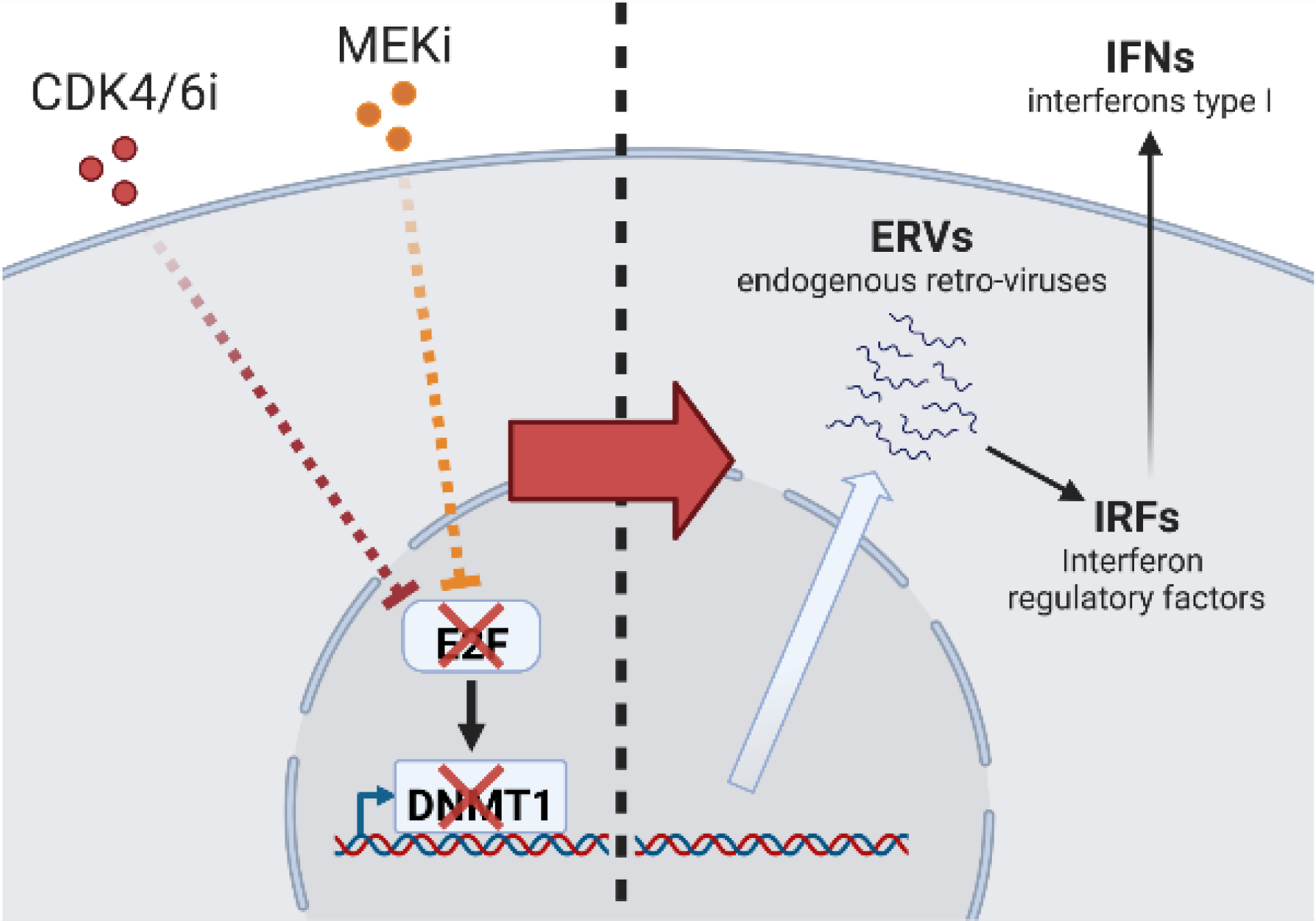
Activation of the viral mimicry upon MEKi and CDK4/6i. MEK and CDK4/6 inhibitors converge on the E2F2 transcription factor. E2F2 regulates the DNA methyltransferase 1 enzyme (DNMT1) responsible for maintaining the DNA methylation profil. The downregulation of DNMT1 leads to a global hypomethylation of the genome and allows the expression of Endogenous retroviruses (ERVs). These transcripts hybridise in the cytoplasm and form dsRNA reminiscent of a viral infection. Cytosolic Pattern Recognition Receptors (PRR) such as RIG-I or MDA5 or endosomal one such as TLR3 then activate the innate immune system leading to an interferon signaling.

**Supplementary Figure 19.**
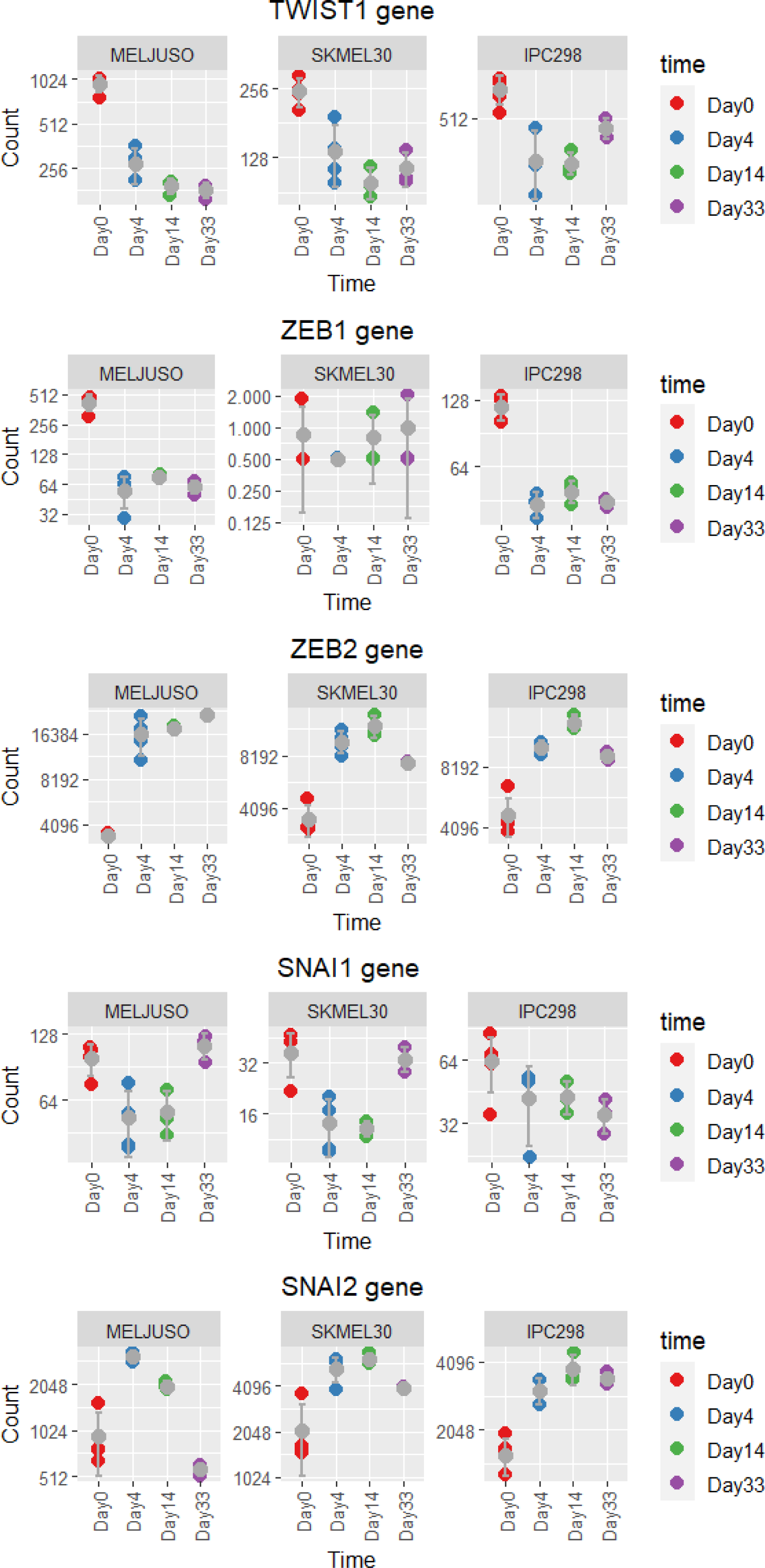
Expression of common regulators of senescence and EMT as listed by (Mir Mohd Faheem et al. 2020) upon MEKi and CDK4/6i.

**Supplementary Figure 20.**
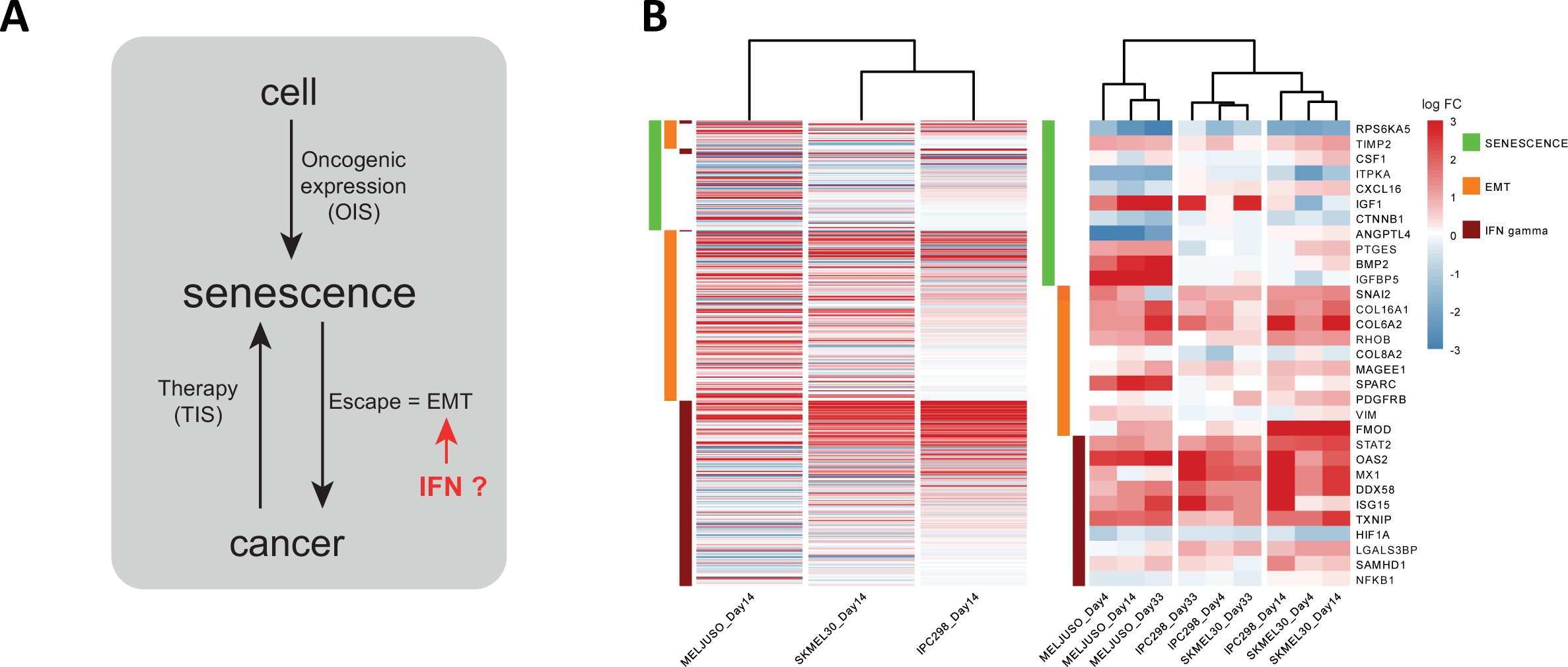
Trend in expression between the “EMT” and IFN gamma gene sets from the hallmarks MSigDB and a recent “senescence” gene list. A. Scheme representing the relationships between the different gene sets. B. Heatmap for the “INTERFERON GAMMA RESPONSE” and “EPITHELIAL MESENCHYMAL TRANSITION” hallmarks gene sets and the “SenMayo” gene list [28].

**Supplementary Figure 21.**
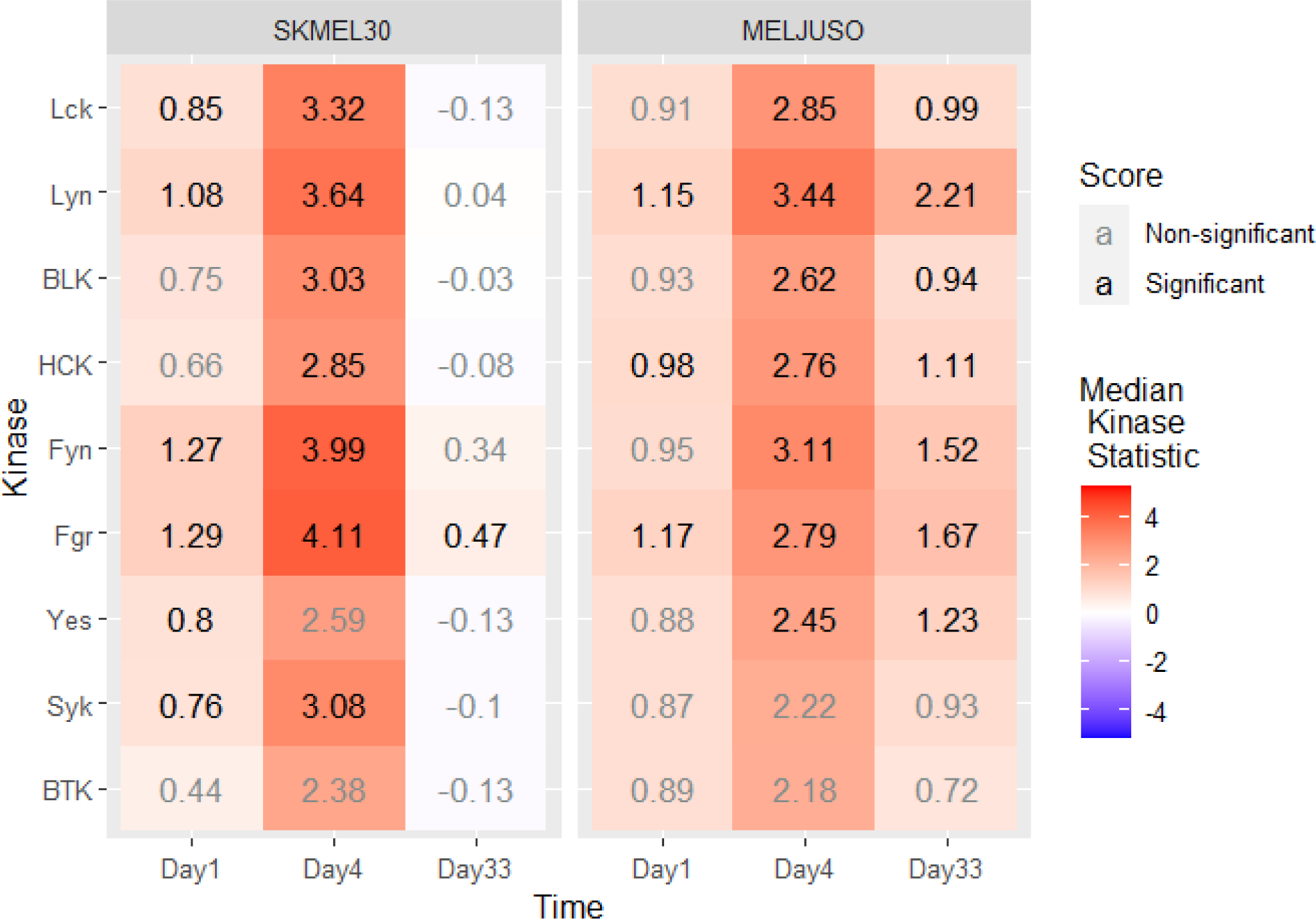
Activation of Src Family Kinases (SFKs). The heatmap represents the “Median Kinase Statistic” for kinases, significant values are represented in black.

**Supplementary Figure 22.**
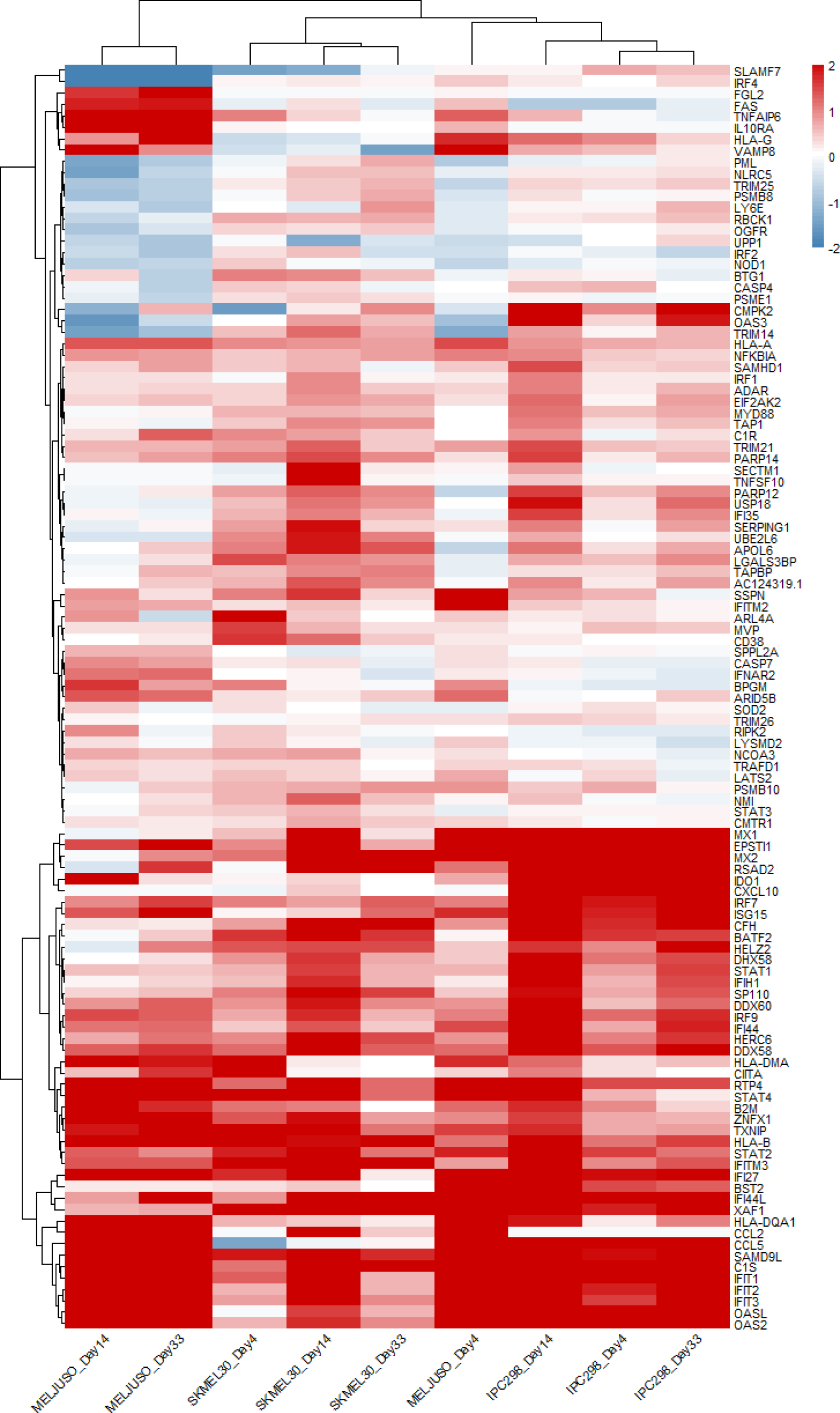
Interferon Gamma response upon MEKi and CDK4/6i. The heatmap represents the logFC of the genes which contribute to the enrichment result. More precisely, it represents a sub-selection of the “INTERFERON GAMMA RESPONSE” gene set which contributes to the “leading edge” (ascendant portion) of the running sum used in GSEA.

**Supplementary Figure 23.**
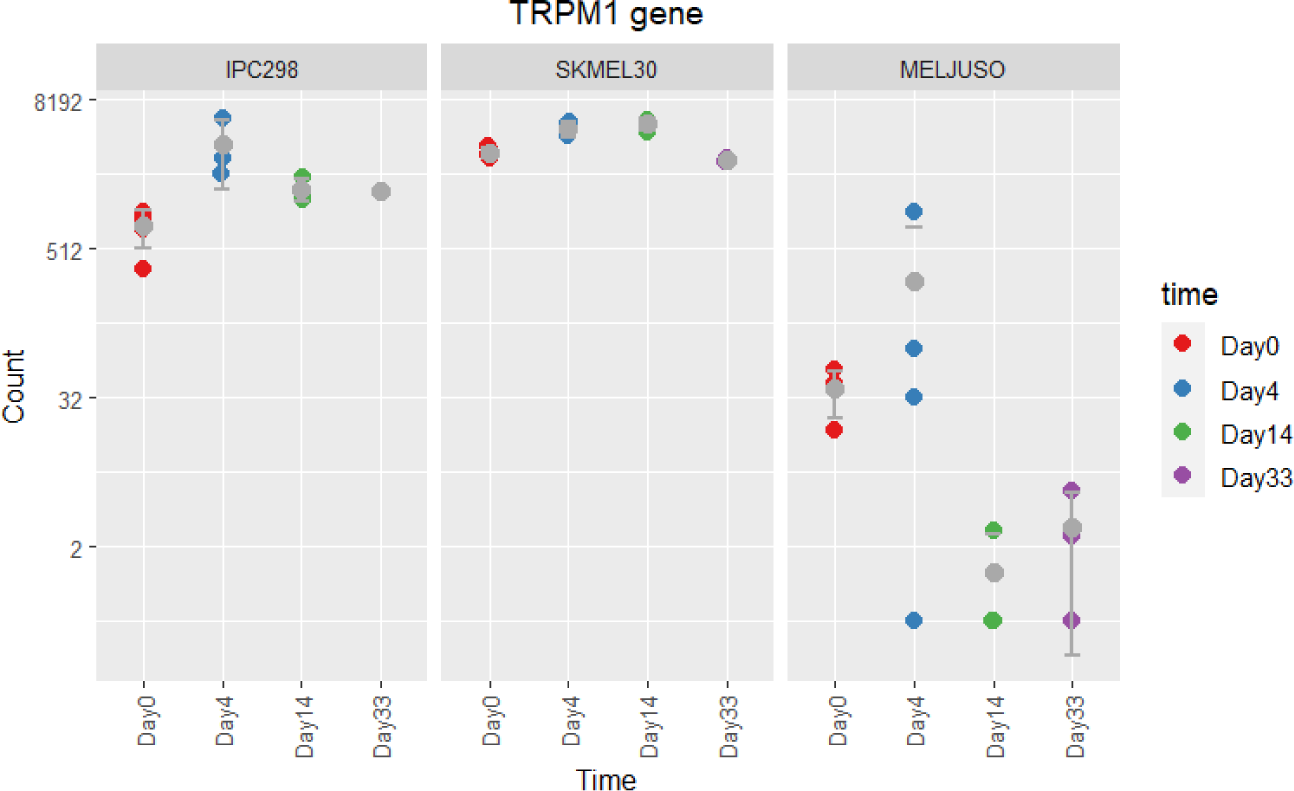
Expression of TRPM1 transcripts in total RNA-seq. TRPM1 transcript contains miR-211-5p in an intronic region.

**Supplementary Figure 24.** Pathway enrichment on qCLASH data using KEGG references. Samples were normalised for read depth and total number of detections for hybrids was summed across technical replicates. Those were further summarised per mRNA and contrasted between conditions. Finally, the top 5% most differentially targeted were used to perform ORA to relate these gene lists to KEGG pathways.

**Supplementary Table 1.**
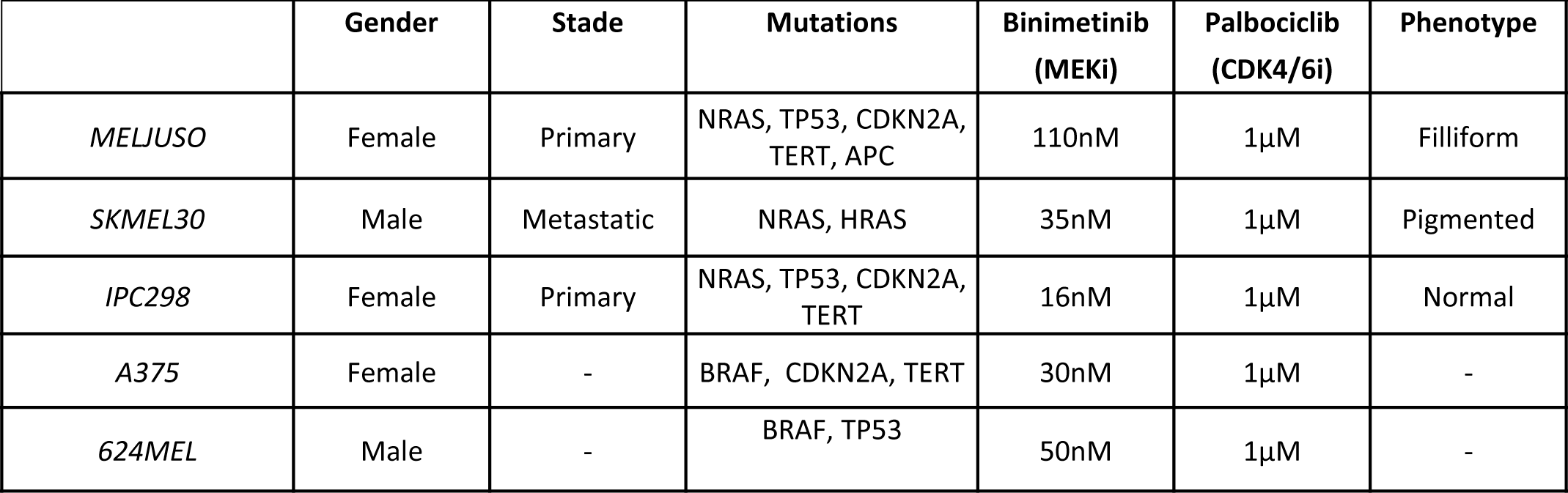
Characteristics and concentrations of inhibitors used for the different cell lines across the study.

|  | Gender | Stade | Mutations | Binimetinib<br>(MEKi) | Palbociclib<br>(CDK4/6i) | Phenotype |
| --- | --- | --- | --- | --- | --- | --- |
| *MELJUSO* | Female | Primary | NRAS, TP53, CDKN2A,<br>TERT, APC | 110nM | 1μM | Filliform |
| *SKMEL30* | Male | Metastatic | NRAS, HRAS | 35nM | 1μM | Pigmented |
| *IPC298* | Female | Primary | NRAS, TP53, CDKN2A,<br>TERT | 16nM | 1μM | Normal |
| *A375* | Female | - | BRAF, CDKN2A, TERT | 30nM | 1μM | - |
| *624MEL* | Male | - | BRAF, TP53 | 50nM | 1μM | - |

**Supplementary Table 2.**
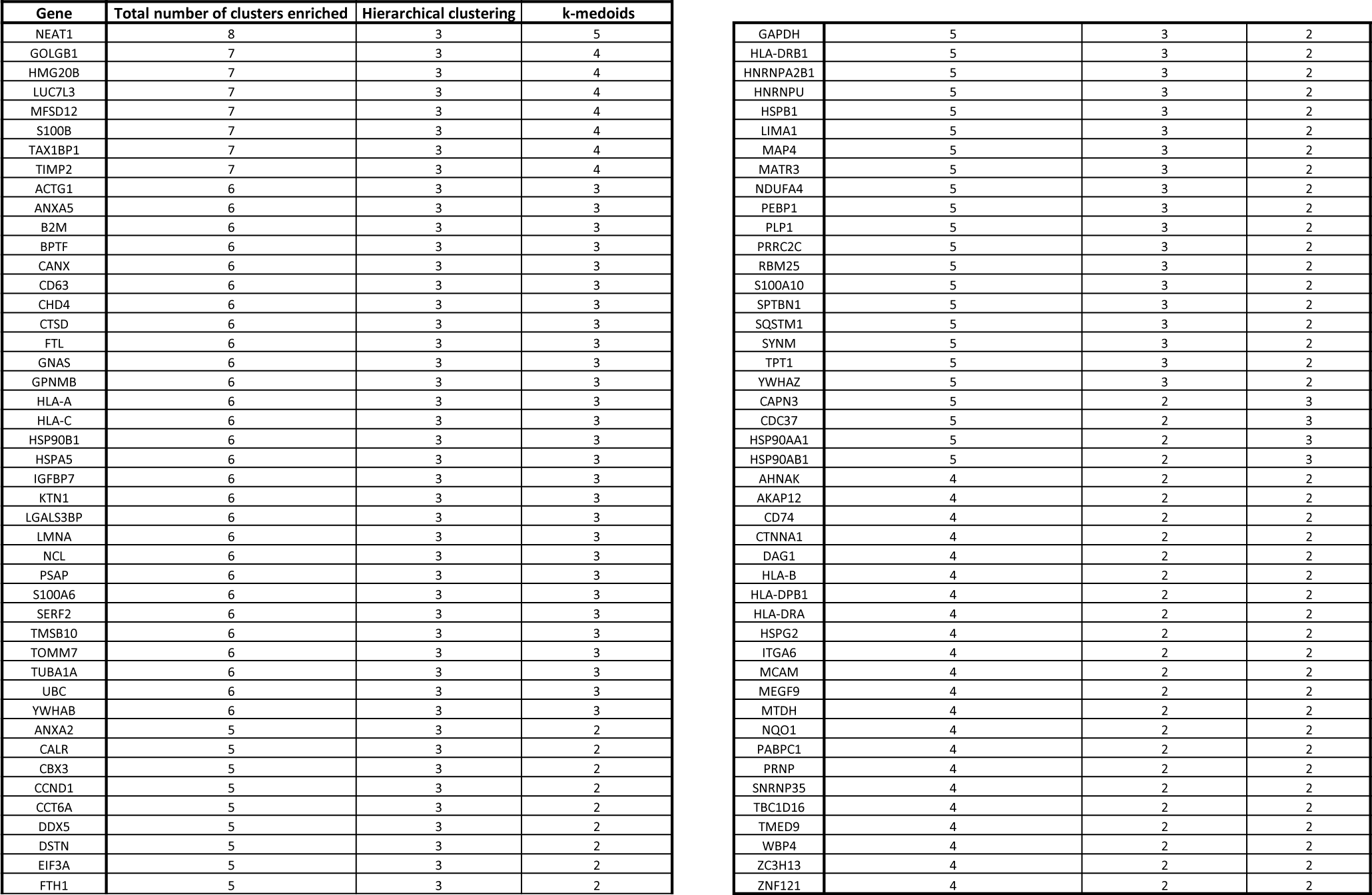
90 genes associated with senescence escape and numbers of clusters enriched for the two algorithms: hierarchical clustering and k-medoids.

**Supplementary Table 3.**
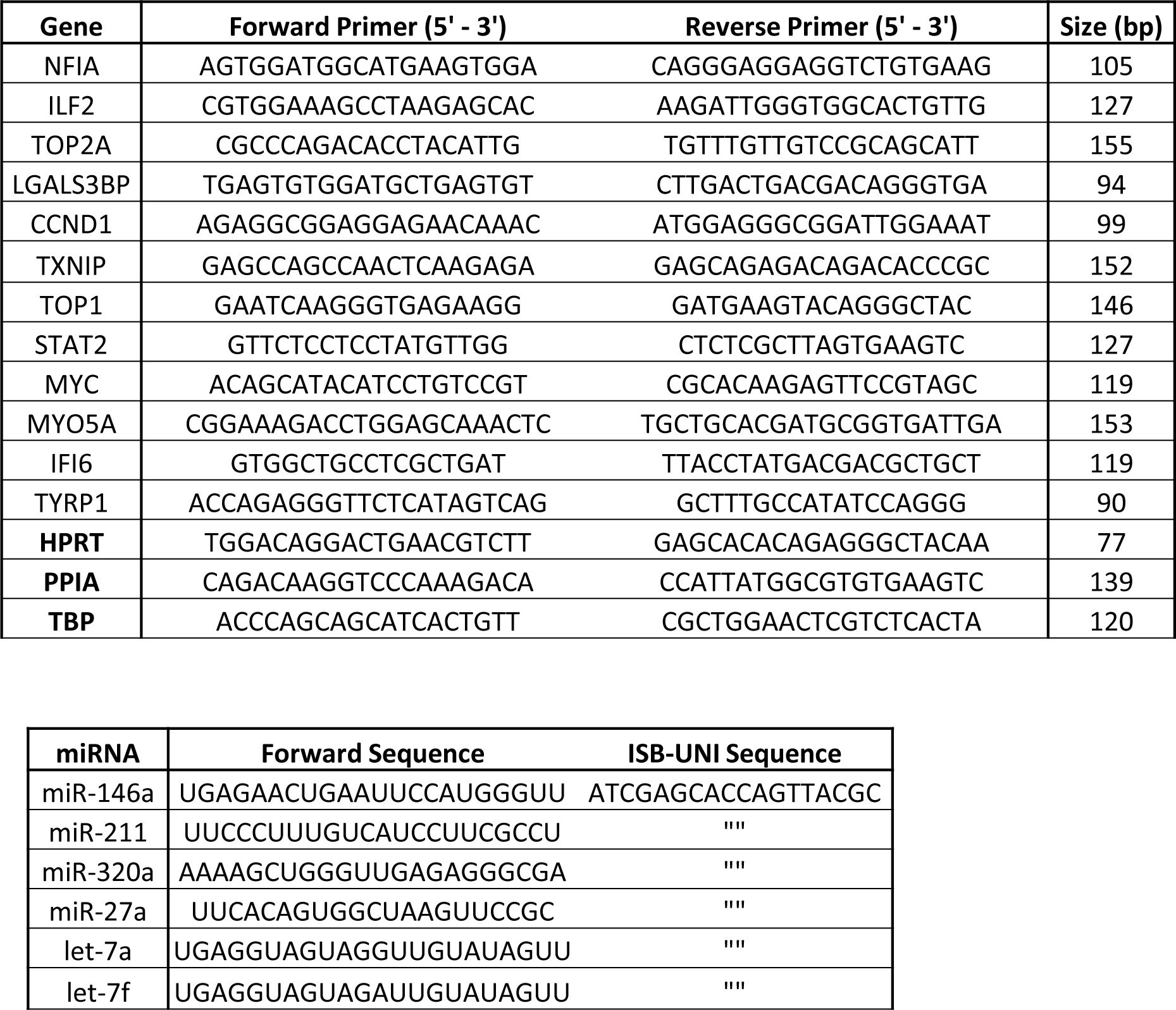
Primer list for mRNA and miRNAs measured by qPCR.

## References

1. Hayflick, L., & Moorhead, P. S. (1961). The serial cultivation of human diploid cell strains. Experimental cell research, 25(3), 585–621.

2. Kumari, R., & Jat, P. (2021). Mechanisms of cellular senescence: cell cycle arrest and senescence associated secretory phenotype. Frontiers in cell and developmental biology, 9, 645593.

3. Ou, H. L., Hoffmann, R., González-López, C., Doherty, G. J., Korkola, J. E., & Muñoz-Espín, D. (2021). Cellular senescence in cancer: From mechanisms to detection. Molecular Oncology, 15(10), 2634–2671.

4. Liao, C., Xiao, Y., & Liu, L. (2020). The dynamic process and its dual effects on tumors of therapy-induced senescence. Cancer Management and Research, 12, 13553.

5. Wang, L., Lankhorst, L., & Bernards, R. (2022). Exploiting senescence for the treatment of cancer. Nature Reviews Cancer, 22(6), 340–355.

6. Campisi, Judith. “Aging, cellular senescence, and cancer.” Annual review of physiology 75 (2013): 685.

7. Campisi, Judith, and Ladislas Robert. “Cell senescence: role in aging and age-related diseases.” Aging 39 (2014): 45–61.

8. Zhu, H., Blake, S., Kusuma, F. K., Pearson, R. B., Kang, J., & Chan, K. T. (2020). Oncogene-induced senescence: From biology to therapy. Mechanisms of ageing and development, 187, 111229.

9. Prieur, A., & Peeper, D. S. (2008). Cellular senescence in vivo: a barrier to tumorigenesis. Current opinion in cell biology, 20(2), 150–155.

10. Roger, L., Tomas, F., & Gire, V. (2021). Mechanisms and regulation of cellular senescence. International Journal of Molecular Sciences, 22(23), 13173.

11. Caksa, Signe, Usman Baqai, and Andrew E. Aplin. “The future of targeted kinase inhibitors in melanoma.” Pharmacology & Therapeutics (2022): 108200.

12. Naeli, Parisa, et al. “The intricate balance between microRNA-induced mRNA decay and translational repression.” The FEBS Journal (2022).

13. Bartel, D. P. (2018). Metazoan micrornas. Cell, 173(1), 20–51.

14. Kozar, I., Philippidou, D., Margue, C., Gay, L. A., Renne, R., & Kreis, S. (2021). Cross-linking ligation and sequencing of hybrids (qCLASH) reveals an unpredicted miRNA Targetome in melanoma cells. Cancers, 13(5), 1096.

15. Malod-Dognin, Noël, et al. “Towards a data-integrated cell.” Nature Communications 10.1 (2019): 1–13.

16. Klein, M. E., Kovatcheva, M., Davis, L. E., Tap, W. D., & Koff, A. (2018). CDK4/6 inhibitors: the mechanism of action may not be as simple as once thought. Cancer cell, 34(1), 9–20.

17. Goel, S., Bergholz, J. S., & Zhao, J. J. (2022). Targeting CDK4 and CDK6 in cancer. Nature Reviews Cancer, 22(6), 356–372.

18. Jochems, F., Thijssen, B., De Conti, G., Jansen, R., Pogacar, Z., Groot, K.,…& Bernards, R. (2021). The Cancer SENESCopedia: A delineation of cancer cell senescence. Cell reports, 36(4), 109441.

19. Laberge, Remi-Martin, et al. “Epithelial-mesenchymal transition induced by senescent fibroblasts.” Cancer Microenvironment 5 (2012): 39–44.

20. Tato-Costa, Joana, et al. “Therapy-induced cellular senescence induces epithelial-to-mesenchymal transition and increases invasiveness in rectal cancer.” Clinical colorectal cancer 15.2 (2016): 170–178.

21. Morales-Valencia, Jorge, et al. “Lipocalin 2 dictates cancer cell plasticity elicited by therapy-induced senescence.” bioRxiv (2022): 2022–03.

22. Rosell, R., Karachaliou, N., Morales-Espinosa, D., Costa, C., Molina, M. A., Sansano, I.,… & Bueno, A.M. (2013). Adaptive resistance to targeted therapies in cancer. Translational lung cancer research, 2(3), 152.

23. Tubita, A., Lombardi, Z., Tusa, I., Lazzeretti, A., Sgrignani, G., Papini, D.,… & Rovida, E. (2021). Inhibition of ERK5 elicits cellular senescence in melanoma via the cyclin-dependent kinase inhibitor p21. Cancer Res.

24. Fields, C. J., Li, L., Hiers, N. M., Li, T., Sheng, P., Huda, T.,…& Xie, M. (2021). Sequencing of Argonaute-bound microRNA/mRNA hybrids reveals regulation of the unfolded protein response by microRNA-320a. PLoS genetics, 17(12), e1009934.

25. Margue, Christiane, et al. “New target genes of MITF-induced microRNA-211 contribute to melanoma cell invasion.” PloS one 8.9 (2013): e73473.

26. Xenos, A., Malod-Dognin, N., Zambrana, C., & Pržulj, N. (2023). Integrated Data Analysis Uncovers New COVID-19 Related Genes and Potential Drug Re-Purposing Candidates. International Journal of Molecular Sciences, 24(2), 1431.

27. Oughtred, R. et al. (2019). The biogrid interaction database: 2019 update. Nucleic Acids Research, 47(D1), D529–D541.

28. Yaveroğlu, Ömer Nebil, et al. “Revealing the hidden language of complex networks.” Scientific reports 4.1 (2014): 1–9.

29. Milenković, Tijana, and Nataša Pržulj. “Uncovering biological network function via graphlet degree signatures.” Cancer Informatics 6 (2008): CIN-S680.

30. Arthur, Liberzon, et al. “The molecular signatures database hallmark gene set collection.” Cell Syst 1 (2015): 417–425.

31. Chan, Michelle, et al. “Novel insights from a multiomics dissection of the Hayflick limit.” Elife 11 (2022): e70283.

32. Lee, Yun Haeng, et al. “Targeting mitochondrial metabolism as a strategy to treat senescence.” Cells 10.11 (2021): 3003.

33. Goel, S., DeCristo, M. J., Watt, A. C., BrinJones, H., Sceneay, J., Li, B. B.,… & Zhao, J. J. (2017). CDK4/6 inhibition triggers anti-tumour immunity. Nature, 548(7668), 471–475.

34. Chiappinelli, K. B., Strissel, P. L., Desrichard, A., Li, H., Henke, C., Akman, B.,… & Strick, R. (2015). Inhibiting DNA methylation causes an interferon response in cancer via dsRNA including endogenous retroviruses. Cell, 162(5), 974–986.

35. Roulois, D., Yau, H. L., Singhania, R., Wang, Y., Danesh, A., Shen, S. Y.,… & De Carvalho, D. D. (2015). DNA-demethylating agents target colorectal cancer cells by inducing viral mimicry by endogenous transcripts. Cell, 162(5), 961–973.

36. Brägelmann, J., Lorenz, C., Borchmann, S., Nishii, K., Wegner, J., Meder, L.,… & Sos, M.L. (2021). MAPK-pathway inhibition mediates inflammatory reprogramming and sensitizes tumors to targeted activation of innate immunity sensor RIG-I. Nature communications, 12(1), 1–15.

37. Huang, Alexander C., and Roberta Zappasodi. “A decade of checkpoint blockade immunotherapy in melanoma: understanding the molecular basis for immune sensitivity and resistance.” Nature immunology 23.5 (2022): 660–670.

38. Takasugi, M., Yoshida, Y., Hara, E., & Ohtani, N. (2022). The role of cellular senescence and SASP in tumour microenvironment. The FEBS Journal.

39. Hu, Xuerui, and Hongqi Zhang. “Doxorubicin-induced cancer cell senescence shows a time delay effect and is inhibited by epithelial-mesenchymal transition (EMT).” Medical science monitor: international medical journal of experimental and clinical research 25 (2019): 3617.

40. Faheem, M. M., Seligson, N. D., Ahmad, S. M., Rasool, R. U., Gandhi, S. G., Bhagat, M., & Goswami, A. (2020). Convergence of therapy-induced senescence (TIS) and EMT in multistep carcinogenesis: current opinions and emerging perspectives. Cell death discovery, 6(1), 1–12.

41. De Blander, Hadrien, et al. “Cellular plasticity: a route to senescence exit and tumorigenesis.” Cancers 13.18 (2021): 4561.

42. Imai, D., Yoshizumi, T., Okano, S., Itoh, S., Ikegami, T., Harada, N.,… & Maehara, Y. (2019). IFN-γ promotes epithelial-mesenchymal transition and the expression of PD-L1 in pancreatic cancer. Journal of Surgical Research, 240, 115–123.

43. Lo, U. G., Bao, J., Cen, J., Yeh, H. C., Luo, J., Tan, W., & Hsieh, J. T. (2019). Interferon-induced IFIT5 promotes epithelial-to-mesenchymal transition leading to renal cancer invasion. American journal of clinical and experimental urology, 7(1), 31.

44. Lee, M., Kim, D. W., Khalmuratova, R., Shin, S. H., Kim, Y. M., Han, D. H.,… & Shin, H.W. (2019). The IFN-γ–p38, ERK kinase axis exacerbates neutrophilic chronic rhinosinusitis by inducing the epithelial-to-mesenchymal transition. Mucosal Immunology, 12(3), 601–611.

45. Saul, Dominik, et al. “A new gene set identifies senescent cells and predicts senescence-associated pathways across tissues.” Nature communications 13.1 (2022): 1–15.

46. Fridman, A. L., & Tainsky, M. A. (2008). Critical pathways in cellular senescence and immortalization revealed by gene expression profiling. Oncogene, 27(46), 5975–5987.

47. Katlinskaya, Y. V., Katlinski, K. V., Yu, Q., Ortiz, A., Beiting, D. P., Brice, A.,… & Fuchs, S.Y. (2016). Suppression of type I interferon signaling overcomes oncogene-induced senescence and mediates melanoma development and progression. Cell reports, 15(1), 171–180.

48. Mullani, N., Porozhan, Y., Mangelinck, A., Rachez, C., Costallat, M., Batsche, E.,… & Muchardt, C. (2021). Reduced RNA turnover as a driver of cellular senescence. Life science alliance, 4(3).

49. Harries, L. W. (2022). Dysregulated RNA processing and metabolism: a new hallmark of ageing and provocation for cellular senescence. The FEBS Journal.

50. Chen, C. H., Hsia, T. C., Yeh, M. H., Chen, T. W., Chen, Y. J., Chen, J. T.,… & Huang, W. C. (2017). MEK inhibitors induce Akt activation and drug resistance by suppressing negative feedback ERK-mediated HER2 phosphorylation at Thr701. Molecular oncology, 11(9), 1273–1287.

51. Ortega-Muelas, M., Roche, O., Fernández-Aroca, D. M., Encinar, J. A., Albandea-Rodríguez, D., Arconada-Luque, E.,… & Sánchez-Prieto, R. (2021). ERK5 signalling pathway is a novel target of sorafenib: Implication in EGF biology. Journal of cellular and molecular medicine, 25(22), 10591–10603.

52. Zhang, J., Pearson, A. J., Sabherwal, N., Telfer, B. A., Ali, N., Kan, K.,… & Tournier, C. (2022). Inhibiting ERK5 overcomes breast cancer resistance to anti-HER2 therapy by targeting the G1–S cell-cycle transition. Cancer Research Communications, 2(3), 131–145.

53. Benito-Jardón, Lucía, et al. “Resistance to MAPK Inhibitors in Melanoma Involves Activation of the IGF1R–MEK5–Erk5 PathwayResistance Mechanisms to MAPK Inhibitors in Melanoma.” Cancer Research 79.9 (2019): 2244–2256.

54. Anerillas, Carlos, et al. “A BDNF-TrkB autocrine loop enhances senescent cell viability.” Nature communications 13.1 (2022): 1–17.

55. Adam, Christian, et al. “Efficient suppression of NRAS-driven melanoma by co-inhibition of ERK1/2 and ERK5 MAPK pathways.” Journal of Investigative Dermatology 140.12 (2020): 2455–2465.

56. Randic, Tijana, et al. “NRAS mutant melanoma: towards better therapies.” Cancer Treatment Reviews 99 (2021): 102238.

57. Cook, Simon J., and Pamela A. Lochhead. “ERK5 Signalling and Resistance to ERK1/2 Pathway Therapeutics: The Path Less Travelled?.” Frontiers in Cell and Developmental Biology 10 (2022).

58. Jiang, W., Cai, F., Xu, H., Lu, Y., Chen, J., Liu, J.,… & Hua, Z. C. (2020). Extracellular signal regulated kinase 5 promotes cell migration, invasion and lung metastasis in a FAK-dependent manner. Protein & cell, 11(11), 825–845.

59. Ortiz, Maria A., et al. “Src family kinases, adaptor proteins and the actin cytoskeleton in epithelial-to-mesenchymal transition.” Cell Communication and Signaling 19.1 (2021): 1–19.

60. Saleiro, D., Blyth, G. T., Kosciuczuk, E. M., Ozark, P. A., Majchrzak-Kita, B., Arslan, A. D.,… & Platanias, L. C. (2018). IFN-γ–inducible antiviral responses require ULK1-mediated activation of MLK3 and ERK5. Science signaling, 11(557), eaap9921.

61. Lee, Bongyong, et al. “MicroRNA-211 modulates the DUSP6-ERK5 signaling axis to promote BRAFV600E-driven melanoma growth in vivo and BRAF/MEK inhibitor resistance.” Journal of Investigative Dermatology 141.2 (2021): 385–394.

62. Santiappillai, Nancy T., et al. “CDK4/6 Inhibition Reprograms Mitochondrial Metabolism in BRAFV600 Melanoma via a p53 Dependent Pathway.” Cancers 13.3 (2021): 524.

63. Teh, Jessica LF, et al. “Metabolic Adaptations to MEK and CDK4/6 Cotargeting in Uveal MelanomaMetabolic Adaptations to Inhibitors in Ocular Melanoma.” Molecular cancer therapeutics 19.8 (2020): 1719–1726.

64. Lee, Jae-Seon, et al. “Targeting oxidative phosphorylation reverses drug resistance in cancer cells by blocking autophagy recycling.” Cells 9.9 (2020): 2013.

65. Corazao-Rozas, Paola, et al. “Mitochondrial oxidative phosphorylation controls cancer cell’s life and death decisions upon exposure to MAPK inhibitors.” Oncotarget 7.26 (2016): 39473.

66. Carpintero-Fernández, Paula, et al. “Genome wide CRISPR/Cas9 screen identifies the coagulation factor IX (F9) as a regulator of senescence.” Cell death & disease 13.2 (2022): 1–13.

67. Martínez-Zamudio, Ricardo Iván, et al. “Escape From Oncogene-Induced Senescence is Controlled by POU2F2 and Memorized by Chromatin Scars.” bioRxiv (2022)

68. Lee, Won Jae, et al. “Genome-wide overexpression screen identifies genes able to bypass p16-mediated senescence in melanoma.” SLAS DISCOVERY: Advancing Life Sciences R&D 22.3 (2017): 298–308.

69. Nagaraj, Karthik, et al. “Long-Term IGF1 Stimulation Leads to Cellular Senescence via Functional Interaction with the Thioredoxin-Interacting Protein, TXNIP.” Cells 11.20 (2022): 3260.]

70. Rambow, Florian, et al. “Toward minimal residual disease-directed therapy in melanoma.” Cell 174.4 (2018): 843–855.

71. Marin-Bejar, Oskar, et al. “Evolutionary predictability of genetic versus nongenetic resistance to anticancer drugs in melanoma.” Cancer Cell 39.8 (2021): 1135–1149.

72. Mölder, Felix, et al. “Sustainable data analysis with Snakemake.” F1000Research 10 (2021).

73. Xiao, Yufei, et al. “A novel significance score for gene selection and ranking.” Bioinformatics 30.6 (2014): 801–807.

74. Metz, Kathleen S., et al. “Coral: Clear and customizable visualization of human kinome data.” Cell systems 7.3 (2018): 347–350.

75. Glaab, E., Baudot, A., Krasnogor, N., Schneider, R., & Valencia, A. (2012). EnrichNet: network-based gene set enrichment analysis. Bioinformatics, 28(18), i451–i457

76. Travis, A. J., Moody, J., Helwak, A., Tollervey, D., & Kudla, G. (2014). Hyb: a bioinformatics pipeline for the analysis of CLASH (crosslinking, ligation and sequencing of hybrids) data. Methods, 65(3), 263–273.

77. Gligorijević, V., Malod-Dognin, N. & Pržulj, N. (2016). Patient-specific data fusion for cancer stratification and personalised treatment. Pac. Symp. Biocomput. 21, 321–332.

78. Ding, C., Li, T., Peng, W. & Park, H. (2006). Orthogonal nonnegative matrix tri-factorizations for clustering. In Proc. ACM SIGKDD International Conference on Knowledge Discovery and Data Mining, Vol. 2006, 126–135, ACM Press.

79. Hart, T., Komori, H. K., LaMere, S., Podshivalova, K., & Salomon, D. R. (2013). Finding the active genes in deep RNA-seq gene expression studies. BMC Genomics, 14(1), 1–7.

80. Pržulj, N. (2019). Analyzing Network Data in Biology and Medicine: An Interdisciplinary Textbook for Biological, Medical and Computational Scientists. Cambridge University Press.

81. Qiao, H. (2015). New svd based initialization strategy for non-negative matrix factorization. Pattern Recognition Letters, 63, 71–77.

82. Pržulj, N., Corneil, D. G. & Jurisica, I. (2004). Modeling interactome: scale-free or geometric? Bioinformatics 20, 3508–3515.

